# Blood-borne immune cells carry low biomass DNA remnants of microbes in patients with colorectal cancer or inflammatory bowel disease

**DOI:** 10.1101/2024.12.16.628609

**Authors:** Y. Morsy, Å. Walberg, P. Wawrzyniak, B. Hubeli, L. Truscello, C. Mamie, A. Niechcial, E. Gueguen, R. Manzini, C. Gottier, S. Lang, S. Scharl, S. Blümel, L. Biedermann, G. Rogler, M. Turina, M. Ramser, H. Petrowsky, I. C. Arnold, S. Zeissig, N. Zamboni, A. Egli, J. H. Niess, P. Hruz, A. Knuth, R. Fritsch, M. G. Manz, M. Wawrzyniak, M. Scharl

## Abstract

The involvement of the human intestinal microbiome in the regulation of immune cell homeostasis, as well as in the pathogenesis of inflammatory bowel disease (IBD) and colorectal cancer (CRC), are well-established^1–4^. Bacteria interact with immune cells at the sites of intestinal inflammation, but also in the CRC tumor microenvironment^1–6^. Indeed, bacterial remnants have recently been detected in human intestinal tissue in patients with IBD, at primary tumor sites and in the metastases of patients with CRC, and in whole blood^3,7,8^. A defective intestinal epithelial barrier is thought to promote bacterial remnant translocation and disease progression^6^. However, it is unknown, how bacterial remnants translocate from the intestine to sites of metastasis or into the circulation. We hypothesized that bacterial remnants translocate within peripheral blood immune cells into the circulation. Here, we thus explored the composition of the detectable microbiome in peripheral blood mononuclear cells (PBMCs) of patients with CRC or IBD compared to healthy controls. The PBMC microbiome profiles partially align with the tumor- and metastasis-derived or intestinal tissue-derived microbiome signatures obtained from the same patients with CRC or IBD, respectively. Our metagenomics data, supported by 16S-rRNA-FISH-Flow, imaging flow cytometry and species-specific qPCR, reveal the presence of translocated bacterial genetic sequences in the patients with CRC and IBD, likely due to an intestinal barrier defect. Pathway and serum metabolomics analysis connected to the metabolic potential of the PBMC-derived microbiome sequences support the onset of microbial translocation in such patients. Thus, our data suggest that in patients with intestinal barrier leakage, such as those with CRC or IBD, there is the potential for the translocation of bacterial remnants into the circulation via peripheral immune cells.

## Introduction

The intestinal microbiome is critical for the maintenance of human health as it is an important contributor to effective epithelial barrier function, immune tolerance, energy production, and digestion, as well as the production of vitamins, bile acids, and short-chain fatty acids^9^. In patients with inflammatory bowel disease (IBD), a reduced diversity and metabolic capacity of the intestinal microbiome has been documented^10–13^. Additionally, patients with active IBD present an enhanced intestinal permeability and increased bacterial DNA load in intestinal tissue compared to healthy controls^14–16^. Thus, it has been hypothesized that the intestinal microbiome affects immune cell homeostasis in patients with IBD and other chronic inflammatory diseases with systemic consequences, either indirectly or directly via the presence of relevant pathogenic bacteria in the periphery, including the circulation^4^. Furthermore, in colorectal cancer (CRC), the intestinal microbiome is found in a dysbiotic state, and bacteria have been found within the tumor microenvironment (TME)^2,3,7,17–20^.

Indeed, several studies have shown the functional implications of such bacterial signatures on cancer progression^1,18,21^. Bacterial genes at primary tumor sites in patients with CRC or oral squamous cell cancer have been observed in cancer and immune cells^3,7^, and regions containing bacterial sequences within the tumor tissue have been associated with an increase in the population of myeloid cells^3^. And, in addition to a detectable microbiome occurring within the primary tumor, it has been shown that CRC-related liver metastases harbor bacteria^7^ and strains of *Fusobacterium nucleatum*, *Bacteroides fragilis,* and *Prevotella spp.* have been identified on a genetic level in both the primary CRC tumor and the corresponding liver metastasis^7^. Animal studies have further revealed that a defective intestinal vascular barrier might promote bacterial translocation, thus promoting cancer metastasis^22^.

The hypothesis that commensal bacteria, or at least bacterial remnants, might translocate from the intestine into the circulation through a potentially defective intestinal epithelial barrier in patients with IBD or CRC is corroborated by the highly sporadic detection of bacterial DNA in the blood of healthy individuals. In an elegant study, *Tan et al.* detected genetic sequences from mainly commensal bacteria derived from the intestinal tract, oral cavity or the genito-urinal tract in whole blood samples of 9,770 healthy individuals. The identified sequences differed from known human pathogens in clinical blood cultures^8^. However, bacterial species were recognized in only 16% of those healthy individuals, and the detected median was one species, suggesting that under homeostatic conditions there is an absence of a consistent core microbiome within human blood^8^. However, the overall microbial DNA load in whole blood in patients with CRC or IBD is higher compared to healthy controls, albeit with yet unclear pathological consequences^27,23^.

As of yet it is unknown how bacteria or bacterial remnants can travel systemically and, in particular, reach sites of distant metastases in patients with CRC or sites of localized inflammatory activity in the case of IBD. But, due to the interaction of bacteria with immune and cancer cells at primary tumor sites^3^, we speculated that such bacterial remnants travel either within cancer cells or immune cells from the primary tumor to sites of metastasis. Here, we show that in patients with CRC or IBD peripheral blood mononuclear cells (PBMCs) may carry translocated bacterial remnants into the circulation, likely due to a defect in their intestinal epithelial barrier.

## Results

### Genetic bacterial sequences are detectable within the PBMCs of patients with CRC or IBD

Firstly, we aimed to explore whether PBMCs might contribute to the translocation of bacterial remnants from sites of primary tumors or from the inflamed intestinal tissue into the blood in patients with CRC or IBD, respectively. We performed metagenomics sequencing on PBMCs derived from 36 patients with CRC, including patients with stages I-III (*n =* 24) and stage IV (*n =* 12), and 56 patients suffering from IBD (ulcerative colitis, UC, *n =* 25; Crohn’s disease, CD, *n =* 31). In addition, we analyzed PBMCs from healthy control individuals (*n* = 20) **(Table S1)**. To exclude potential contamination affecting the results, we used multiple lab reagent controls **(Table S2)**. We performed extensive bioinformatics data cleaning procedures according to recommended guidelines to avoid potential pitfalls of low-biomass microbial analyses and to recognize the reagent microbiome in our samples **(Fig. S1A-C, Table S3, Methods)**^24,25^. Further, we only selected species for further analysis that are well-known to colonize the human gut and are thus ecologically plausible. After performing each step of this data decontamination pipeline, 208 bacterial species (126 in tissue only and 82 in both PBMCs and tissue) were detectable at a rate of at least 10 specific reads in at least 3 patients for at least one disease in either tissue and/or PBMCs **(Fig. S1A-C, Table S4**).We found a higher microbial richness as measured by Chao1 in PBMCs from patients with metastatic and non-metastatic CRC compared to PBMCs from patients with IBD **(Fig. 1A**, **Table S5)**. Beta diversity analysis based on Bray-Curtis presented by Principal Coordinates Analysis (PCoA) demonstrated a cluster of PBMCs from patients with non-metastatic CRC were separated from patients with IBD and healthy control individuals. These results were statistically supported by PERMANOVA analysis **(Fig. 1B, Table S6)**. Notably, while based on Chao1 alpha diversity, there was no clear correlation between age and the number of bacterial species detected within the PBMCs of younger and older individuals (Fig. S2A-G).

**Figure 1.**
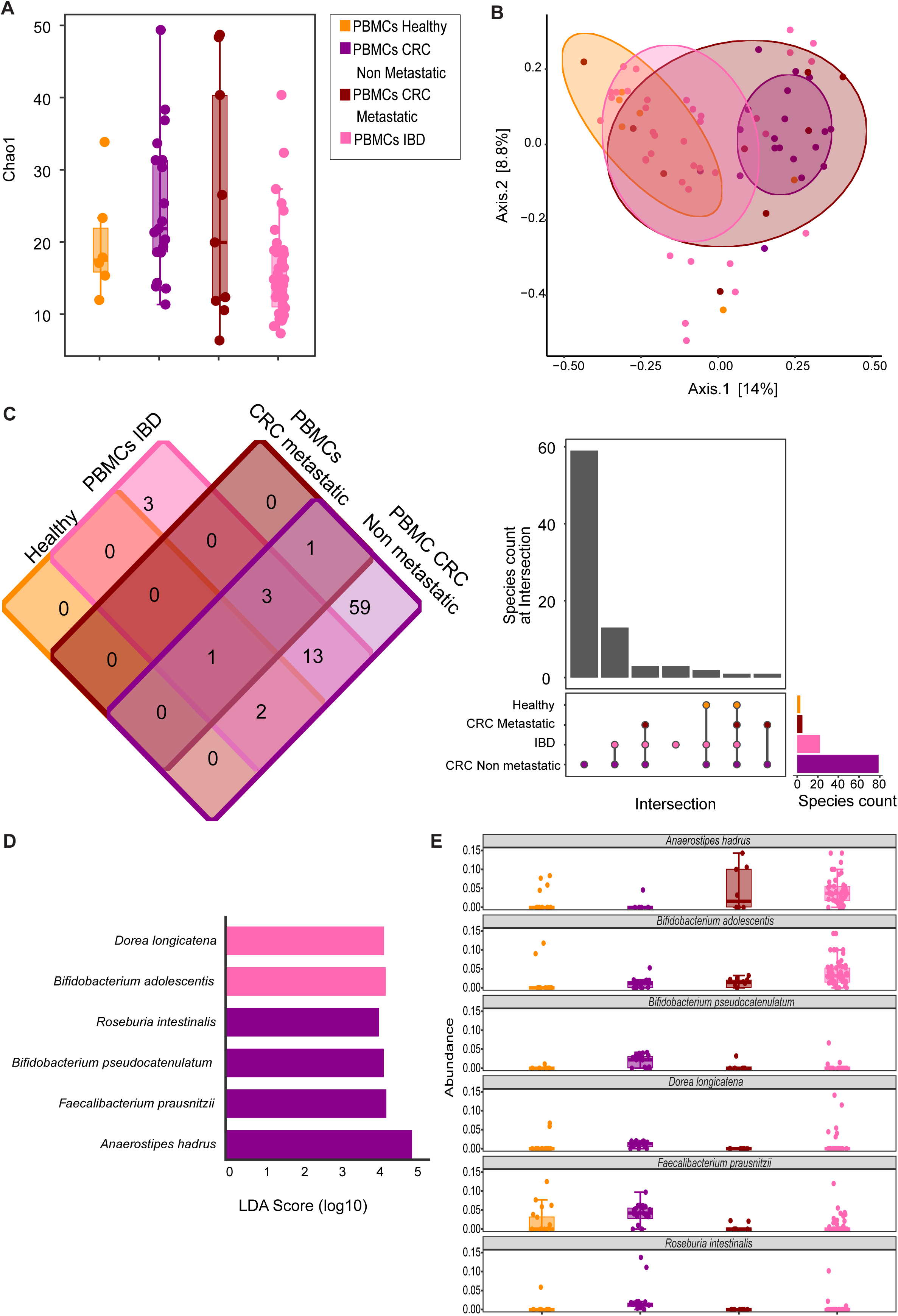
The bacterial profile of PBMCs in a cohort of patients with CRC, IBD, and healthy controls. Specific samples were removed due to rarefaction, which was conducted to normalize the results. **(A)** Chao1 alpha diversity index diversity measurements of bacterial taxa were detected in PBMCs from patients with IBD (*n* = 37), CRC non-metastatic (*n* = 19), metastatic CRC (*n* = 9), as well as in healthy controls (*n* = 6). Specific samples were removed due to rarefaction normalization. **(B)** Beta diversity analysis based on Bray-Curtis presented by Principal Coordinates Analysis (PCoA) showed differences between PBMCs from healthy individuals and PBMCs from patients with CRC and IBD. **(C)** A Venn diagram and an upset plot show the number of common bacterial species in different diseases. **(D)** Effect size analysis (LDA score, log10) shows different taxa detected in PBMCs from each group, as in A, with an LDA score of more than 3. (**E**) Relative abundance (normalized to a total of 1) of the six species with LDA score above 3.

After applying our strict filtering and decontamination criteria, we still detected bacterial DNA in the PBMCs from all patients and all healthy controls. To check the commonly detected bacterial DNA in PBMCs between different diseases, we set up a threshold of 10 reads to consider the bacterial DNA as present in each condition **(Fig. S1D)**. Therefore, PBMCs from patients with non-metastatic CRC had the highest number of diverse bacterial species (*n =* 79), followed by IBD (*n =* 22). In contrast, PBMCs from patients with metastatic CRC (*n =* 5) exhibited only very few bacterial species, and the lowest number of species was detected in PBMCs from healthy individuals (*n =* 3) **(Fig. 1C, Table S4)**. Of the three species detected in healthy individuals in our samples, *Anaerobutyricum hallii* and *Haemophilus parainfluenzae* had previously been reported in a comprehensive analysis of the whole-blood microbiome of 9770 healthy individuals^8^, which included extensive filtering for contaminants.

Out of the 82 species detected in PBMCs overall, 31 exhibited a remarkable difference between at least two groups **(Fig. S3**). Further, we could define disease-specific PBMC patterns as determined by the effect size (LDA score, log10, **Fig. 1D-E)**. *Roseburia intestinalis, Bifidobacterium pseudocatenulatum, Faecalibacterium prausnitzii* and *Anaerostipes hadrus* were distinctive for patients with non-metastatic CRC, while *Bifidobacterium adolescentis* and *Dorea longicatena* were indicative for PBMCs from patients with IBD **(Fig. 1D-E**). Notably, *Anaerostipes hadrus* featured a high LDA score for non-metastatic CRC as it was enriched in PBMCs from patients with metastatic CRC or with IBD.

### Bacterial DNA is present in specific PBMC subsets from patients with CRC and IBD

Next, we aimed to confirm the presence of bacterial DNA in PBMCs and to determine the specific immune cell subsets carrying bacterial DNA in the peripheral circulation. We integrated the detection of the bacterial 16S rRNA using a fluorescence *in situ* hybridization (FISH) probe into our flow cytometry staining of PBMCs (16S rRNA-FISH-Flow) to further validate our findings **(Figs. 2A-B, Figs. S4-7)**. Confirming our previous finding of a very low number of bacterial species detected in PBMCs from healthy controls using metagenomic sequencing, we detected almost no bacterial 16S rRNA signal in PBMCs from healthy controls (*n =* 9) using 16s rRNA FISH Flow. In contrast, we identified the presence of bacterial 16S rRNA in selective immune cell subtypes of PBMCs from patients with CRC (*n =* 10), particularly in CD4^+^ T cells and in CD3^−^CD14^+^ monocytes. When analyzing the PBMCs from patients with IBD *(n =* 9), we detected bacterial 16S rRNA to an overall lesser extent and primarily within the CD3^−^CD14^+^ monocytes. To further validate our findings, we expanded our analysis and adopted the imaging flow cytometry technique. This method enabled us to visualize cells containing the 16S rRNA gene while simultaneously exhibiting characteristic immune cell markers. In this manner, we confirmed our metagenomics findings and validated the presence of 16S rRNA within specific immune cell populations at a single-cell level **(Figs. 2C-D).**

**Figure 2.**
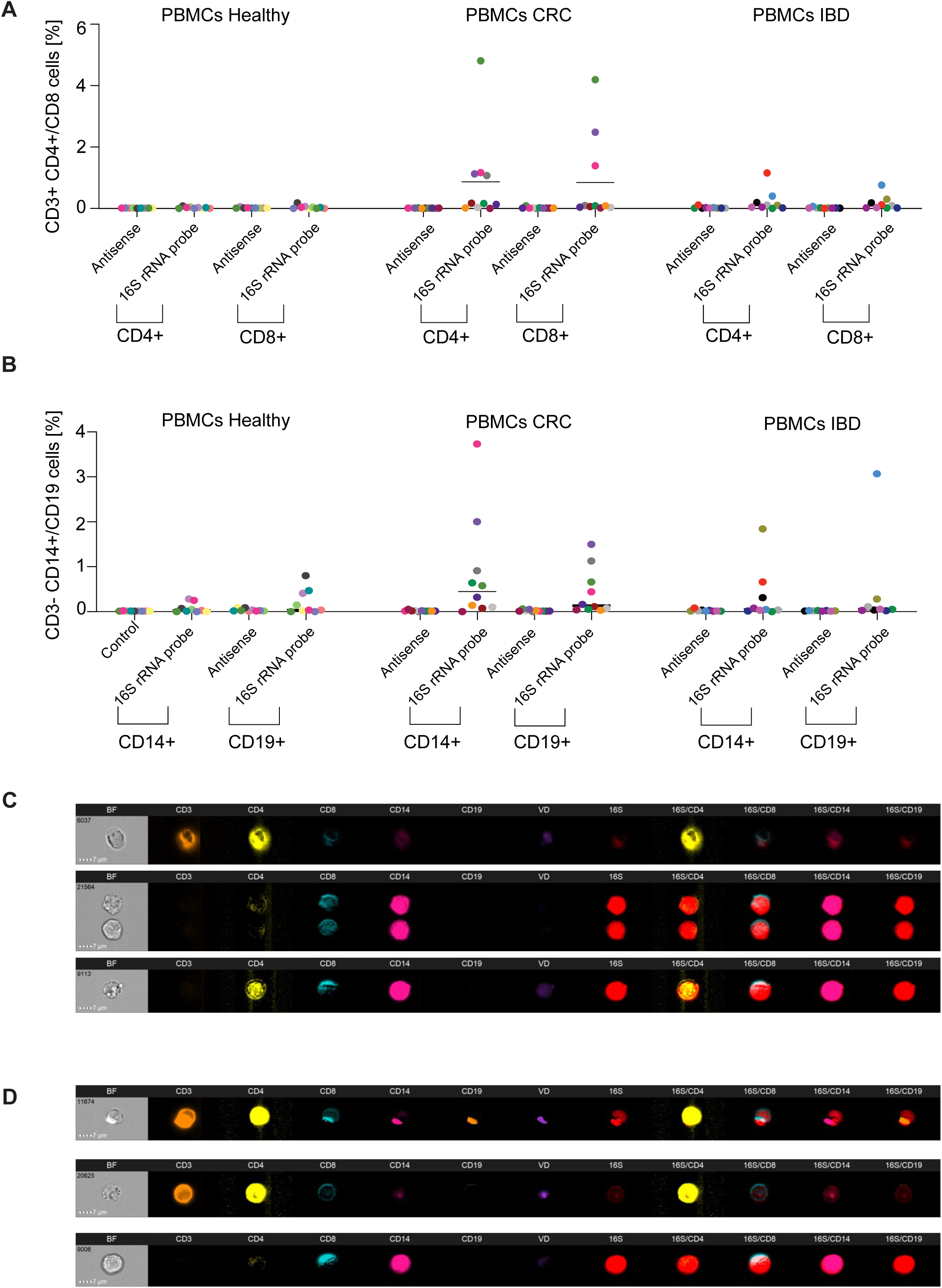
Detection of 16S rRNA-positive cells in PBMCs from patients with CRC or IBD. PBMCs from healthy controls, *n* = 9, CRC patients, *n* = 10 and IBD patients, *n* = 9 were stained for surface markers CD3, CD4, CD8, CD14 and CD19, and 16S rRNA probe and complementary antisense as a control. (**A**) Detection of 16S rRNA probe in CD3^+^CD4^+^ T cells and CD3^+^CD8^+^ T cells in comparison to antisense probe. (**B**) Detection of 16S rRNA probe in CD3^−^CD14^+^ monocytes and CD3^−^CD19^+^ B cells in comparison to antisense probe. Each symbol represents one donor. (**C**) Representative pictures of PBMCs from CRC donor 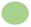 positive for 16S rRNA probe together with CD3, CD4, CD8 and CD14 cell markers visualized by Imaging Flow Cytometry. (**D**) Representative pictures of PBMCs from IBD donor 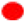positive for 16S rRNA probe together with CD3, CD4, CD8 and CD14 cell markers visualized by Imaging Flow Cytometry. Data is shown as means +/− SD, Two-way ANOVA test followed by Bonferroni’s multiple comparison test was performed.

### The bacterial sequences detectable within PBMCs and tumor-derived tissue from patients with CRC suggest translocation of bacterial remnants from the intestine

Having demonstrated the presence of bacterial remnants in PBMCs of patients with CRC and IBD, we next aimed to identify the potential origin of the bacterial species found in the PBMCs. Firstly, we explored patients with CRC from whom either PBMCs, primary tumor, and adjacent colon tissue were available (*n = 16* non-metastatic, *n =* 4 metastatic) and patients with CRC from whom PBMCs, metastatic liver and adjacent liver tissue were available (*n =* 5) **(Table S1).** We observed a markedly higher alpha diversity as measured by Chao1, meaning a higher bacterial richness, in both unaffected colon tissue and tumor tissue from patients with non-metastatic CRC compared to their matching PBMCs **(Fig. 3A)**. Beta diversity analysis using Bray-Curtis revealed remarkable differences between PBMCs from patients with CRC and matching unaffected colon and primary tumor tissues **(Fig. 3B)**. Notably, *Anaerostipes hadrus* and *Collinsella aerofaciens* were the most dominant bacterial species in PBMCs from patients with CRC. These species were also detectable in the respective matched intestinal and metastatic CRC tissue, as well as the respective adjacent intestinal and liver tissue (**Fig. 3C**). In contrast, we found that *Haemophilus parainfluenzae, Prevotella sp. E2-28* and *Clostridioides difficile* were the most abundant species in the PBMCs from patients with liver metastasis and their corresponding CRC liver metastatic tissue. Of note, these species were less abundant in the PBMCs from patients with non-metastatic CRC, and they were almost absent in their matching intestinal CRC and adjacent tissue (**Fig. 3C**). In contrast, *Bacteroides fragilis* and *Phocaeicola vulgatus* were highly detectable in intestinal CRC and intestinal adjacent tissue, but mostly absent in the PBMCs and the liver tissues. *Fusobacterium nucleatum* was mainly detectable in the intestinal tumor tissue from patients with non-metastatic CRC, but also in the PBMCs of those patients (**Fig. 3C)**. These patterns suggest distinct translocation profiles of specific bacterial species from the intestine into the PBMCs in patients with CRC. Overall, for 82 species we were able to demonstrate remarkable differences in relative abundance between the cancer tissue and their corresponding adjacent tissue compared to the PBMCs from the same patients **(Fig. S8)**.

**Figure 3.**
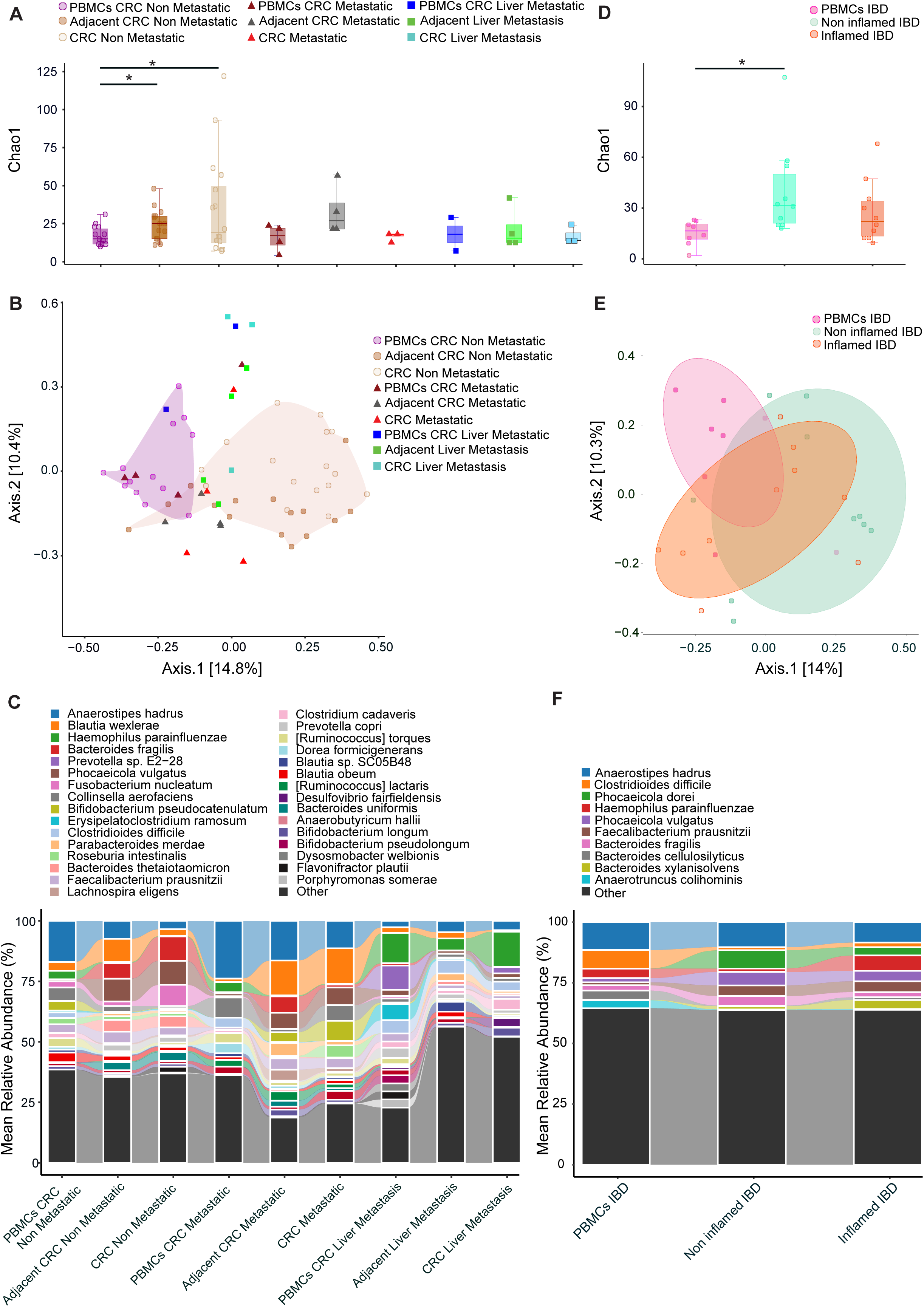
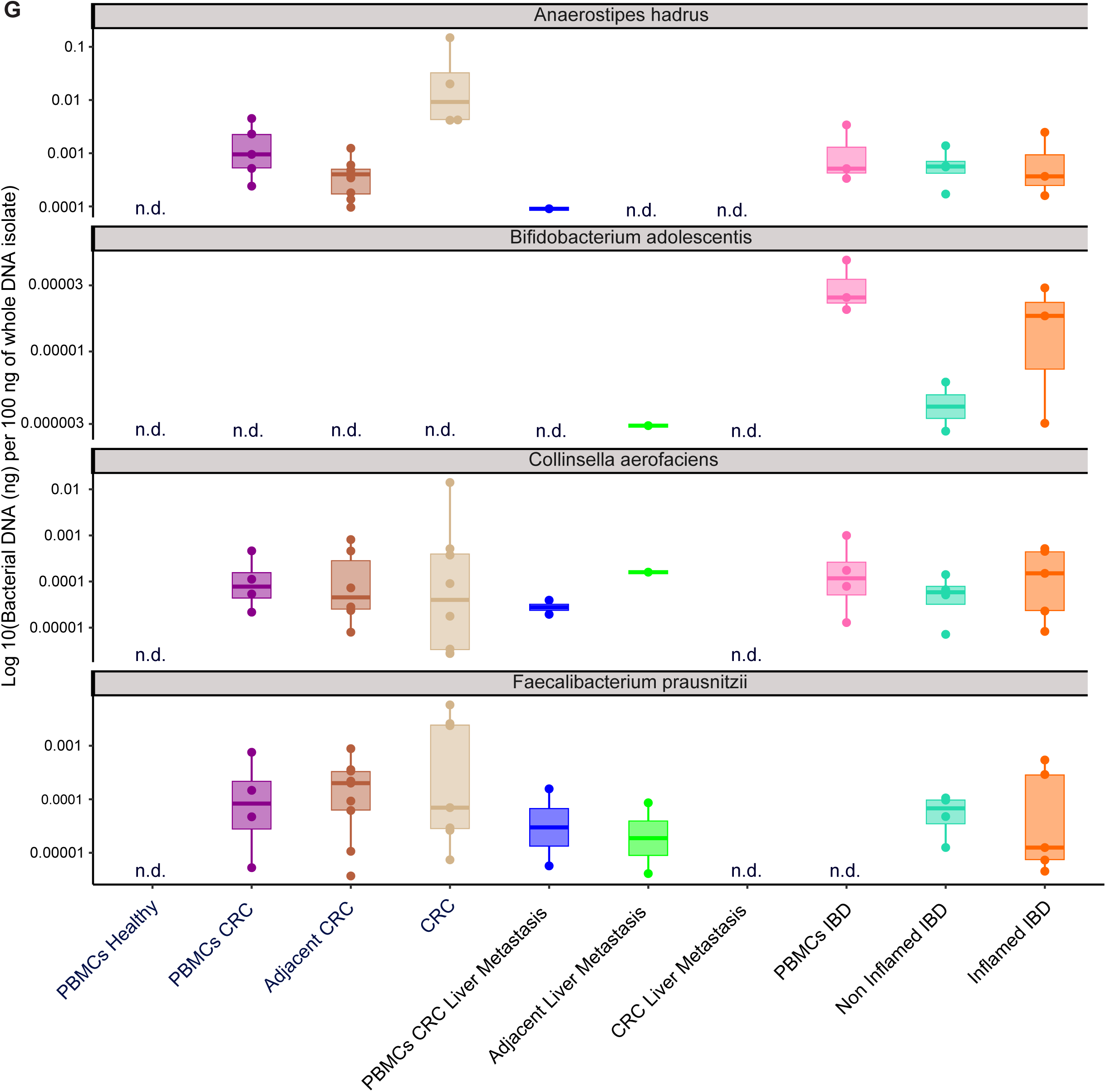
Bacterial profiles of PBMCs and tissues from a cohort of patients with CRC or IBD. Specific samples were removed due to rarefaction, which was conducted to normalize the results. **(A)** Chao1 diversity index analysis between PBMCs (n = 20 and 8 samples excluded), corresponding unaffected colon tissue and CRC tissue from the same patients (*n* = 20 and 5 samples excluded), PBMCs from patients with metastatic CRC (*n* = 4 and one sample excluded), and PBMCs from patients with CRC liver metastasis, corresponding unaffected liver tissue and CRC liver metastatic lesion from matched patients (*n* = 5 and 3 samples excluded). **(B)** Beta diversity as assessed by Bray Curtis analysis. **(C)** Mean relative abundance of most abundant bacterial species pattern across PBMCs and different CRC tissue. **(D)** Chao1 diversity index analysis between PBMCs (*n* = 11 and 3 samples excluded), corresponding non-inflamed colon tissue, and inflamed tissue from the same patients (*n* = 11 and one sample excluded). **(E)** Beta diversity assessed using the Bray Curtis analysis. **(F)** Relative abundance of most abundant bacterial species pattern across PBMCs and different colon tissue from patients with IBD. **(G)** Detection of *Anaerostipes hadrus*, *Bifidobacterium adolescentis*, *Collinsella aerofaciens* and *Faecalibacterium prausnitzii* with qPCR on whole DNA isolates from PBMCs from CRC patients (n = 7, in at least 4/7 patients, DNA was detectable, except for *Bifidobacterium adolescentis*), adjacent CRC (*n* = 13, in least 6/13 patients, DNA was detectable, except for *Bifidobacterium adolescentis*), tumor tissue from CRC patients (*n* = 10, in at least 4/10 patients, DNA was detectable, except for *Bifidobacterium adolescentis*), PBMCs from patients with CRC liver metastasis (*n* = 4, in at least 1/4 patients, DNA was detectable, except for *Bifidobacterium adolescentis*), adjacent liver tissue from patients with CRC liver metastasis (*n* = 4, in at least 1/4 patients, DNA was detectable, except for *Anaerostipes hadrus*), liver metastasis tissue from patients with CRC liver metastasis (*n* = 4, in all 4 patients, DNA was n.d.), PBMCs from IBD patients (*n* = 8, in at least 3/8 patients, DNA was detectable, except for *Faecalibacterium prausnitzii*), non-inflamed IBD colon tissue (*n* = 8, in at least 2/8 patients DNA was detectable) and inflamed IBD colon tissue (*n* = 9, in at least 3/9 patients, DNA was detectable) and PBMCs from healthy controls (*n =* 4, in all 4 patients, DNA was n.d.). Data are presented as log10 transformed predicted bacterial DNA concentrations in ng, based on standard curve values generated from pure bacterial DNA of the respective species (Fig. S10). n.d. = not detectable.

### The bacterial profile of PBMCs and intestinal tissue from patients with IBD suggests the translocation of bacterial remnants from the intestine

Next, we aimed to first confirm in patients with IBD our finding in the PBMCs from patients with CRC, and, secondly, we wanted to explore whether the PBMC-derived microbial sequences in patients with IBD might also potentially originate from the intestine. We analyzed a cohort of 11 patients with IBD from which PBMCs, as well as non-inflamed and inflamed intestinal tissue samples, were available **(Table S1)**. By studying alpha diversity as measured by Chao1, we identified a significant difference between non-inflamed intestinal tissue and PBMCs from the same patients with IBD. The bacterial richness was lowest in the PBMCs from these patients, suggesting the presence of more dominant species in such PBMCs than in the tissue (**Fig. 3D**). By a beta diversity analysis using Bray-Curtis we revealed a clear clustering between the PBMCs from patients with IBD and non-inflamed tissue from the same patients, while there was again no clear difference between PBMCs and inflamed tissue from the same patients (**Fig. 3E**).

As in CRC, we observed that *Anaerostipes hadrus* was the most abundant species in PBMCs from patients with IBD and it was also highly abundant in their inflamed and non-inflamed intestinal tissue (**Fig. 3F**). While less abundant overall, *Phocaeicola vulgatus*, *Faecalibacterium prausnitzii,* and *Bacteroides fragilis* were highly abundant in both inflamed and non-inflamed intestinal tissues, and they were found in lower relative abundance in the PBMCs. In contrast, *Haemophilus parainfluenzae* and *Clostridioides difficile* were highly abundant in the PBMCs and the inflamed tissue, but less detectable in the non-inflamed intestinal tissue. In contrast, *Anaerotruncus colihominins* was almost exclusively present in the PBMCs, while *Phocaeicola dorei* and *Bacteroides xylanisolvens* was almost exclusively found in the intestinal tissue (**Fig. 3F**).

To further confirm our findings, we next performed species-specific qPCR on the whole DNA samples that were already used for the metagenomics sequencing. However, we did not have sufficient samples left over to determine the presence of bacterial species in paired PBMCs and respective matched tissue samples. Nevertheless, we confirmed the specific presence of *Anaerostipes hadrus*, *Collinsella aerofaciens*, *Faecalibacterium prausnitzii*, and *Bifidobacterium adolescentis* in PBMCs and tissues from patients with CRC and IBD, respectively **(Fig. 3G, Fig. S9+10, Table S7)**. These data further support the notion that the translocation of remnants of intestinal bacteria into the PBMCs occur in patients with an underlying intestinal barrier defect.

### A different pattern of bacterial remnants is detectable in PBMCs and intestinal tissue or tumor tissue between patients with CRC and IBD

When comparing all CRC and IBD samples, alpha diversity analysis using the Chao1 metric indicated variations in microbial diversity across different groups **(Figs. 3A+D)**. Notably, PBMCs from patients with metastatic CRC exhibited a relatively high alpha diversity compared to patients with non-metastatic CRC or IBD. We observed marked differences in the abundance of specific microbial species among the groups (**Figs. 4A+B**). For instance, *Prevotella sp. E2-28* and *Fusobacterium nucleatum* were highly abundant in tissue samples from patients with CRC. Conversely, *Vescimonas coprocola* and *Simiaoa sunii* showed a low abundance in these tissues and were almost absent in the PBMCs. Notably, species such as *Faecalibacterium sp. L3839* and *[Ruminococcus] lactaris* were highly abundant in metastatic tissues. The most abundant species, however, including *Phocaecola vulgatus* and *Anaerostipes hadrus*, showed a varied distribution across the groups, indicating their ubiquitous presence.

**Figure 4.**
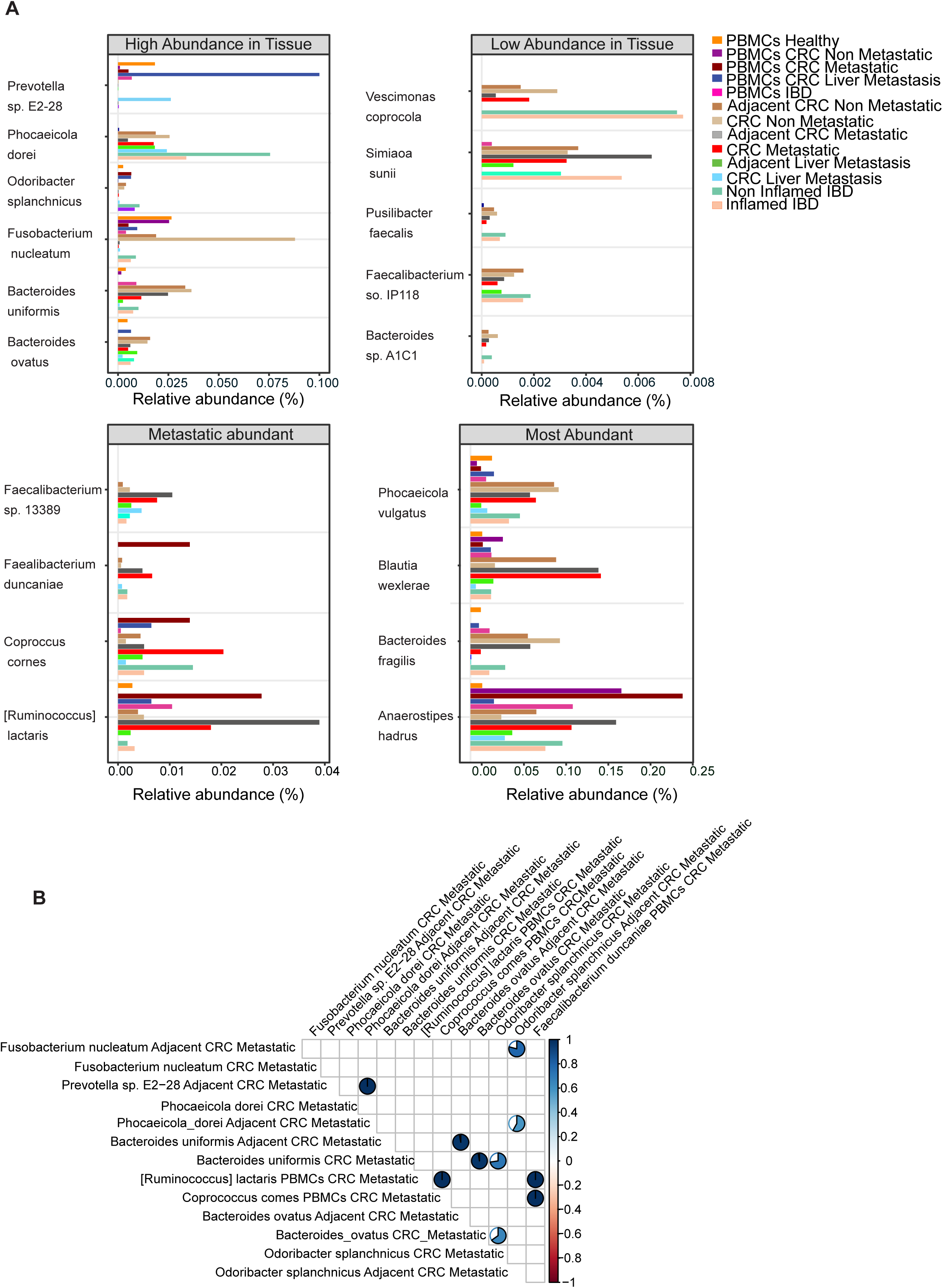
Comparison of bacterial species detected in patients with CRC or IBD in contrast to PBMCs from healthy controls. **(A)** Histograms show the relative abundance (%) of bacterial species categorized as high tissue abundant (> 0.02%), low tissue abundant (< 0.02%), metastatic abundant (> 0.02%), and most abundant (> 0.02% in both tissue and PBMC samples), with each bar representing the mean relative abundance for each species across different sample types. **(B)** Correlation heatmap of bacterial species represented in Fig 4A between samples from CRC metastatic tissue and PMBCs. Only significant correlations with p-values less than 0.05 and Spearman correlation above 0.5 are represented by a pie diagram, and blank squares are insignificant.

### Metabolite profiling and metagenomics sequencing analysis support the functional relevance of the PBMC-microbiome interaction in patients with CRC and IBD

Next, we aimed to further delineate the potential functional relevance of the PBMC-derived microbiome in patients with CRC and IBD. We applied untargeted metabolomics to profile serum metabolites accounting for the metabolic potential of the detected microbes. We obtained serum from 28 patients with CRC (stages I-III *n = 19*, stage IV *n = 9*), 12 patients with IBD, and 10 healthy control individuals, all with a previously analyzed PBMC microbiome. We observed distinct differences in the metabolite profile of patients with CRC and IBD compared to healthy controls **(Fig. 5A)**. A total of 635 metabolites were significantly different between the three groups, and differentially regulated metabolites showed an apparent clustering in the heat-map **(Fig. S11)**. We noted differences for 445 metabolites between patients with IBD and healthy individuals, for 539 metabolites between patients with CRC and healthy individuals, and for 80 metabolites between patients with IBD and CRC. Metabolites altered in either patients with CRC or IBD were used in Metabolite Set Enrichment Analysis (MESA) and pathway analysis, highlighting key metabolic pathways affected in these two patient groups. Here, we observed a disease-specific enrichment, or at least partially, of specific classes of metabolites, including isothiocyanates, diazines, pyrimidine nucleotides, lactones, and benzenes **(Fig. 5B)**. Pathway enrichment analysis indicated that most of the altered metabolites were involved in phenylalanine, tyrosine, and tryptophan biosynthesis, linoleic acid metabolism, histidine, starch, and sucrose metabolism **(Fig. 5C)**. Next, we performed a correlation network analysis between altered serum metabolites in patients with CRC and IBD and pathways from corresponding PBMC metagenomics functional analysis data. This analysis revealed that in particular ketogenesis was highly positively correlated, and that plasmalogen degradation was clearly negatively correlated, with the detected serum metabolites (**Figs. 5D-E, Tables S 8-9)**.

**Figure 5.**
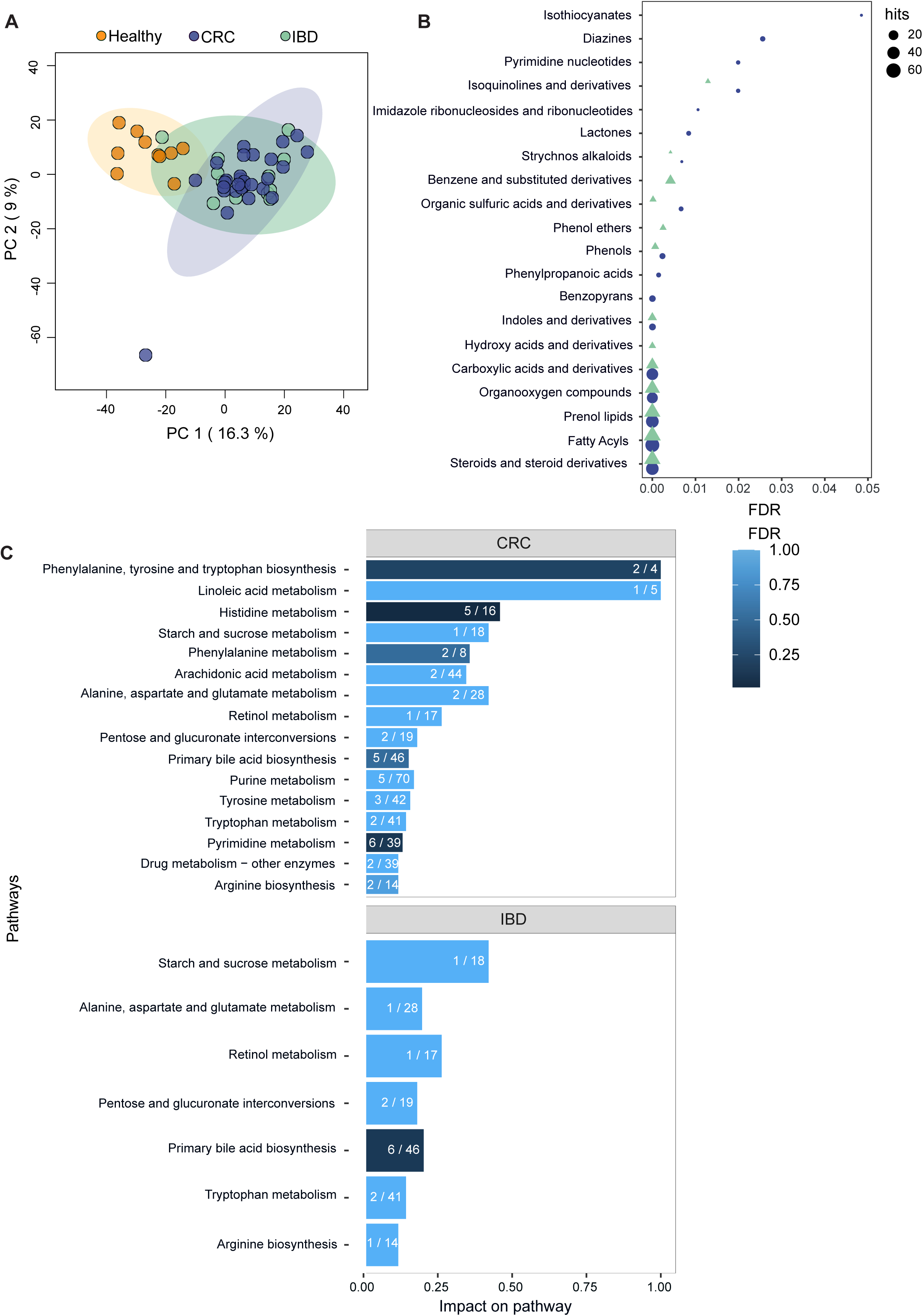
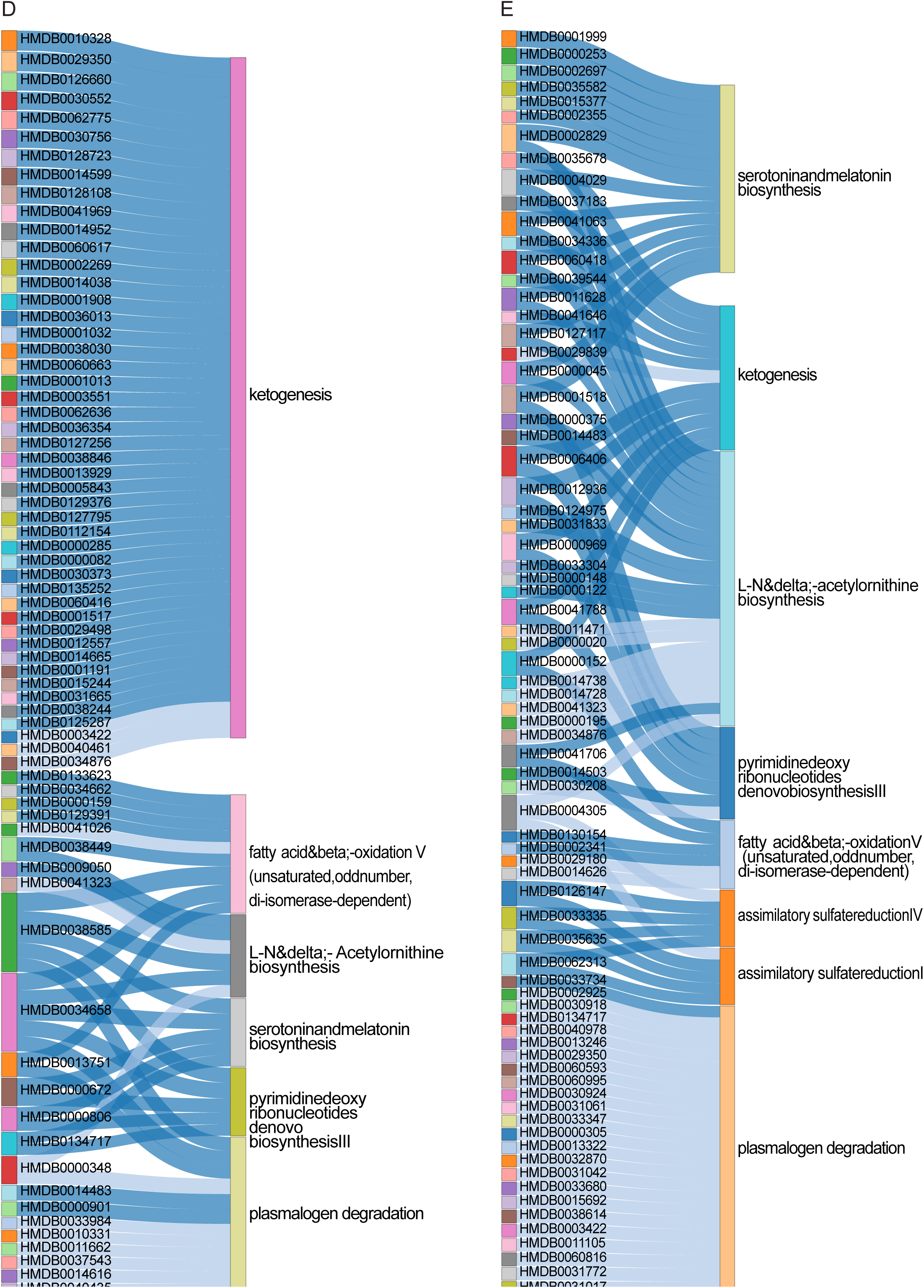
The serum metabolite profile of a cohort of patients with CRC or IBD and in healthy controls. **(A)** Principal component analysis (PCA) of serum metabolites in patients with CRC (*n =* 28), IBD (*n* = 12), and healthy controls (*n =* 10). Each dot represents one patient. **(B)** Metabolites set enrichment analysis of 524 common altered metabolites compared to healthy. **(C)** Pathway analysis of the detected metabolites. **(D**) Correlation network between metabolites detected in serum samples from patients with CRC non-metastatic and altered pathways from metagenomics functional analysis of PBMC samples from patients with CRC non-metastatic. **(E)** Correlation network between metabolites detected in serum samples and altered pathways resulted from metagenomics functional analysis of PBMC samples from patients with IBD. Only significant correlations with p-values less than 0.05 and Spearman correlation above 0.5 are represented by dark blue lines and less than −0.5 by light blue lines.

## Discussion

Here, we provide evidence for a genomic bacterial pattern present within circulating immune cells in the peripheral blood of patients with CRC and IBD. Our data further suggest that the PBMC-derived microbial sequences are likely the result of bacterial translocation from the intestine, particularly in patients with an underlying defect in the intestinal epithelial barrier, which commonly occurs in CRC and IBD.

The role of the microbiome in the pathogenesis of chronic inflammatory and malignant diseases is becoming increasingly evident^1–4^. Recently, it has been demonstrated that primary tumors, as well as metastasis, from patients with CRC exhibit bacterial genetic sequences^3,7^; however, the means by which these bacterial remnants might be transported from the primary tumor to the sites of metastasis, *e*.*g*., in the liver, is still unclear. Our data support the concept that in patients with CRC, PBMCs might serve as carriers of bacterial remnants from the intestine and the primary tumor, respectively, towards the sites of metastasis. This data is also in line with the known pro-metastatic roles of certain immune cells recruited to the tumors^26^.

Our data is of interest because, to date, the presence of bacteria in human immune cells has only been described in the context of infectious diseases. Pathogenic *Salmonella* or *Mycobacteria* species can invade and proliferate within dendritic cells and/or macrophages^27,28^. Further, in mouse models of obesity, live bacteria have been shown to translocate from the intestine to adipose tissue via the microbial pattern receptor CD14^29^. In CRC and oral squamous cell cancer bacterial sequences have been detected intracellularly in both the cancer cells and the immune cells of the primary tumors^3,7^. In patients with CRC, *Fusobacterium nucleatum* has been detected in the primary tumor tissue in the intestine, as well as in the liver metastasis^7^ and was additionally suggested to reach tumors via the blood^30^. To date, however, the presence of bacteria or bacterial remnants has not been described in circulating PBMCs from patients with CRC or from those with IBD. Thus, this may be a potential means by which bacterial remnants are transported to and end up in peripheral tissues and metastases.

We should note that we observed differences between the microbiome of PBMCs from patients with CRC and IBD. The detected differences in bacterial relative abundances in samples of the same cell type, processed at a single sequencing center, analyzed with the identical pipeline and following the recommendations for reagent microbiome detection, provide substantial evidence that these dissimilarities indeed originate from biological variations and not from recently described technical limitations^24,25,31^. To support our observation, we further confirmed the detection of bacterial sequences in PBMCs from patients with CRC and IBD using a well-established 16S rRNA FISH probe (EUB338)^32,33^. Thus, our data demonstrate at least the presence of translocated bacterial genetic sequences, likely bacterial remnants, within the PBMCs of patients with CRC or IBD. Thus, the observed presence of the detected bacterial sequences (regardless whether this indicates the presence of live or dead bacteria) might have an impact on immune cell function and thus on disease pathogenesis, which would be crucial to investigate in future studies.

We also recognized a higher species count for PBMCs from cases of non-metastatic CRC *vs.* those from cases of metastatic CRC, which at first glance seems counter-intuitive within the context of our bacterial translocation model as it would suggest an increased barrier impairment as the adenocarcinoma progresses would lead to potentially greater bacterial translocation. One potential explanation might be, however, that patients with metastatic CRC may have a compromised immune function due to advanced disease or treatments like chemotherapy, which could impact bacterial translocation or its recognition by the immune system^34^. Additionally, changes in the gut microbiota composition towards an even further reduced diversity as the disease progresses might affect the diversity of bacteria available for translocation and thus result in reduced diversity even in the PBMCs, suggesting the presence of fewer, but more dominant, species than in the PBMCs of patients with early-staged CRC^35^. This hypothesis is reflected in our data where in PBMCs from patients with liver metastasis there is a high abundance of bacteria, such as *Haemophilus parainfluenzae*, *Prevotella sp. E2−28*, *Erysipelatoclostridium ramosum* or *Clostridioides difficile*, which are clearly less abundant in PBMCs of patients with early-staged CRC.

This concept of reduced diversity might also account for the observed overall trend that younger patients exhibited a higher Chao1 diversity within their PBMCs than older patients. This was especially pronounced in healthy individuals, as well as in patients with metastatic CRC, and indeed a somewhat surprising observation as one might expect that the intestinal barrier is more effective in young people than in elderly people^36^. As an explanation, one might speculate that the highly diverse intestinal microbiome being present within the intestine of younger people might allow for sporadically more diverse bacterial remnants to pass through the intestinal barrier than in older patients. The microbiome of elderly patients is characterized by clearly less diversity and thus likely more dominant, and also potentially more pathogenic, bacterial species^37^. Thus, such more dominant bacterial species might translocate primarily through the corrupted intestinal barrier into the PBMCs, leading to the greater detection of more dominant species within the PBMCs of the elderly participants in our study. Such a mechanism could also occur in patients with CRC and liver metastasis, where we indeed found higher abundances of dominant, species, such as *Haemophilus parainfluenzae*, *Prevotella sp. E2−28* or *Clostridioides difficile*. Thus, the presence and translocation of such dominant species would overrule the effect of a disrupted barrier in elderly people. In patients with IBD the effect of a pronounced dysbiosis in elderly patients compared to younger patients might be somewhat limited as IBD *per se* is characterized by a heavy dysbiosis independent from the patient’s age^38^. However, we should note that our cohort was very small to address such differences on a statistically relevant level and further and extensive research beyond the scope of this study would certainly be required to address this issue in detail.

In our metagenomics data analysis we have strictly adhered to the guidelines for data decontamination in low-biomass microbial studies and paid also particular attention to the recognition of the reagent microbiome in our samples^24,25^. In this way, we aimed to ensure that we did not face recently described problems in certain, mostly cancer-related, microbiome studies^31^. Thus, having performed such stringent contamination removal from our data, our data are now in contrast to previous literature in the field. Notably, many of the previous studies reporting a microbiome within the blood or the tissue of various tumors did not perform such rigorous data decontamination, nor did they report findings that are problematic from an ecological perspective and/or exhibit a high abundance of non-human commensal *Proteobacteria* and finally also commonly contradicting themselves, as, for example, discussed by Ghihawi *et al.*^31^. In particular, some of those publications report a broad amount of *Proteobacteria*, *e*.*g*., *Sphingomonadaceae* or *Caulobacteraceae*, which are typical environmental and not human commensal bacteria^31^. Such reported overrepresentation of environmental *Proteobacteria* in the blood or tumor tissue is most likely due to insufficient/incomplete data decontamination and thus recognition of a reagent microbiome, thus denying a common blood microbiome. It also does not make sense from a biological/ecological perspective, given that *Bacteroidetes*, *Firmicutes* and *Actinobacteria* are the main phyla found in the human intestine^39^ and the source of detected bacterial remnants in human blood/PBMCs is likely the intestine in individuals with an impaired intestinal epithelial barrier. Further, additional questions arise regarding the validity of such findings in many of the previous studies as bacterial sequences had even been described in cancers of sterile tissues.

We should also note that we detected some uncommon immune cell marker combinations, *e*.*g*., CD3^−^CD4^+^ cells, CD4^+^CD8^+^ double-positive cells, and also CD8^+^CD14^+^ double-positive cells. However, those marker combinations have particularly been demonstrated to play a role in immune responses to bacterial antigens and/or the immune response in patients with cancer or chronic inflammatory diseases^40–45^. We detected only a limited number of sequences from bacterial species that are well-known to reside within the human intestine, which further contradicts the notion that the detection of the bacterial sequences by our metagenomics analysis is a result of erroneously assigned human reads in metagenomics sequencing data^31^. Hence, collectively, our data indeed support the presence of at least bacterial remnants in blood cell compartments^8,23,46^.

We have previously demonstrated that some of those identified bacteria, *e*.g*., Roseburia intestinalis* or *Faecalibacterium prausnitzii*, indeed affect anti-tumor immunity *in vivo*^17^. *Anaerostipes hadrus,* which proved to be highly detectable in both the PBMCs and intestinal tissue from patients with CRC, was identified in a recent study as a critical modulator of the metabolism of 5-fluorouracil, which is a state-of-the-art chemotherapy for the treatment of CRC. This suggests an important role for this bacterium in patients with CRC^47^. All three of these bacteria have been shown to be highly reduced in the feces of patients with CRC^17^, but are present here at high levels in the primary tumor tissue and, at least on a genetic level, in the PBMCs of those patients. A further validation of our data is the fact that we identified *Fusobacterium nucleatum* at high levels in the primary tumor tissue of patients with CRC, but not in the intestine of patients with IBD, and at low levels again in PBMCs from patients with CRC and CRC liver metastatic tissue.

By focusing on the distinct bacterial species present in the intestine-derived tissue, as well as the corresponding PBMCs, from patients with IBD and CRC, respectively, we found characteristic patterns for a relevant number of species that might be closely related to their respective functionality. In the intestinal tissue from patients with IBD and CRC, but not in their PBMCs, we detected a high abundance of *Ruminococci*; *e*.*g*., the mucolytic *Ruminococcus torques*, *Bacteroides fragilis* or *B. uniformis*. Such mucus degradation might aggravate the defect in the intestinal barrier, allowing additional bacteria to penetrate deeper tissue layers. Correspondingly, we detected a high abundance of various *Lachnospiaraceae*, such as *Anaerostipes hadrus, Blautia wexlerae* or *Roseburia intestinalis,* and *Faecalibacteria*, in the intestinal tissue, but also in the PBMCs of those patients. Notably, in PBMCs from patients with a severe intestinal barrier defect; *i*.*e*., patients with active IBD and metastatic CRC, we detected high abundances of pathogenic bacteria, such as *Clostridioides difficile* or *Haemophilus parainfluenzae*. These observations are in line with the recent identification of systemic anti-microbiota IgG repertoires, indicating bacterial translocation from the intestine in patients with IBD^48^.

The key novelty of our work is the identification of bacterial sequences in PBMCs of patients with CRC and IBD, as previous studies only focused on the microbiome of fecal, tissue or whole-blood samples, but not on peripheral immune cells. Though our study is limited by the lack of matching primary CRC tissue and corresponding CRC liver metastasis tissue from the same patients due to clinical limitations. Nevertheless, our data demonstrate the presence of a limited number of bacterial sequences that are common to the colonic primary tumor, the PBMCs from patients with CRC and the metastatic CRC liver lesions. Additionally, the overlap of bacterial species found in unaffected normal colon tissue, in CRC tissue, inflamed IBD tissue, and in PBMCs from patients with CRC and IBD, respectively, potentially suggests a translocation of bacterial remnants, possibly due to increased permeability of the gut barrier as described for CRC and IBD^49–51^. A further limitation that could not yet be addressed in our present study is whether the detected bacterial genetic sequences in PBMCs might be derived from live bacteria.

Overall, we provide evidence here for the presence of bacterial remnants in circulating immune cells of patients with CRC and IBD. The comparison between PBMCs and the corresponding tissue samples from the same patients identified some common bacterial species, indicating potential mutual microbial signatures, suggesting that a bacterial translocation from the intestine might be the source of the microbial sequences detected in the PBMCs of patients with CRC or IBD (**Figure S12**). Our findings contribute to the understanding of the complex pathogenesis of CRC and IBD and raise the possibility that the presence of microbial remnants in PBMCs might exert a potential mechanistic role in the pathogenesis of such diseases.

## Acknowledgments

We thank Damina Balmer for providing editorial assistance. Flow cytometry was performed with support of the Flow Cytometry Facility, University of Zurich.

This work was supported by the Stiftung Experimentelle Biomedizin (MS), Swiss National Science Foundation grant No 320030_184753 (MS), Swiss National Science Foundation grant No 320030E_190969 (MS), The Swiss Cancer League grant no. KFS-5372-08-2021-R (MS), Research grant from the Fondazione San Salvatore (MS), Research grant from the Stiftung für wissenschaftliche Forschung an der Universität Zürich grant No STWF-22-009 (MS), Research grant from the ISREC Foundation (MS, ICA), Lighthouse project grant of the Comprehensive Cancer Center Zurich (CCCZ) und University Medicine Zurich (UMZH) (MS), Wilhelm-Sander Foundation grant no. 2021.104.1 (MS), Research grant from the Iten-Kohaut Foundation (BH), Research grant from the University of Zürich “Forschungskredit” (BH), Research grant from Novartis Foundation for medical-biological Research (BH), Research grant from the Fond’Action contre le cancer (BH). SB was supported by a Walter and Gertrud Siegenthaler Fellowship, a grant from the Hartmann Mueller Foundation and a fellowship from the Fonds zur Foerderung des Akademischen Nachwuchses, University of Zurich.

## Author contributions

Conceptualization: YM, PW, MW, ÅW, MS

Methodology: YM, PW, AN, LT, ÅW, BH, RM, SS, CG, SL, LB, NZ, MS

Investigation: ÅW, PW, BH, LT, CM, AN, EG, RM, CG, SL, SB, GR, MT, MR, HP, ICA, NZ, SZ, AE, JHN, PH, AK, RF, MGM, YM, MS

Visualization: YM, ÅW, PW

Funding acquisition: MS, BH, ICA, SB

Project administration: MS

Supervision: BH, MGM, MS

Writing – original draft: YM, ÅW, MS

Writing – review & editing: All authors

## Inclusion & ethics statement

All collaborators listed as co-authors in this study fulfill all authorship criteria required by Nature Portfolio. This work included local researchers throughout the research process and has been determined in collaboration with local partners. Roles and responsibilities were agreed amongst collaborators ahead of the research. This research was not severely restricted or prohibited in the settings of the researchers. The research projects have been approved by the local ethics review committees and local environmental protection and biorisk-related regulation were sufficient. This work did not result in stigmatization, incrimination, discrimination or other personal risks to participants. Local and regional research relevant for this study was taken into account in citations.

## Data and materials availability

Sequencing data (fastq) have been deposited at NCBI Home - BioProject - NCBI (nih.gov) or at the European Nucleotide Archive (ENA). They are publicly available as of the publication date under the accession number PRJNA1024674. Metabolomics data has been deposited at MassIVE and is publicly available as of the date of publication under accession number MSV000095692.

## Supplementary Figures

**Fig S1: The different steps describing our data decontamination pipeline.**

**(A)** Filtering criteria with the corresponding number of species left after each filtering step. **(B)** Hierarchical correlation to detect the possible contamination clusters. **(C)** The principal coordinate analysis is based on weighted unifrac to check possible batch effect. **(D)** Read counts of the detected species in PMBCs of different health conditions where species are listed on the Y axis, and the read counts on the X axis. Each dot represents the corresponding read count in a single patient.

**Fig S2: Overview of Demographics, Diversity, and PBMC Profiles by Disease and Age Group**

Demographic distributions, microbial diversity, and PBMC profiles across different disease categories. **(A)** The graph presents the gender distribution among healthy, CRC, and IBD patients highlighting comparable gender ratios in each group. **(B)** Graph demonstrates the regression analysis between the age and Chao1 index. **(C)** Boxplot illustrates the age distribution by disease, showing median ages and interquartile ranges. **(D)** The density plots of age distributions across the three disease groups visualizes age-related trends. **(E)** Graph categorizes age distributions further by disease and age group (younger < 43 and older ≥ 43). **(F)** Graph demonstrates microbial alpha diversity (Chao1 index) against age for the studied cohorts. **(G)** Visualization of PBMC profiles using PCoA-differentiated disease categories and age groups.

**Fig S3: Representation of statistically significant species in the PBMCs.**

The heatmap represents the relative abundance of statistically significant species across PBMCs and their prevalence in the patient samples. Black squares represent significance in the corresponding comparison. Kruskal-Wallis test followed by pairwise Wilcoxon tests with Benjamini-Hochberg adjusted p-value less than 0.05, is considered significant.

**Fig S4: Bacterial DNA detection in PBMCs infected with *E. coli*.**

**(A)** Flow cytometry plots on live PBMCs treated with antibiotics (gentamicin (100 ug/ml), doxycycline (5 ug/ml)) and infected with different concentrations (MOI 0, MOI 0,5, MOI 1, MOI 2) of *Escherichia coli* (*E.coli*). Positive cells for 16s rRNA were calculated based on negative control complementary antisense. **(B)** Summary plot of PBMCs showing increase of detection of 16s rRNA positive cells with increased concentration of E. coli, *n* = 3-9. **(C)** Flow cytometry plots representing 16S rRNA positive cells in CD4^+^, CD8^+^ T cells, CD14^+^ monocytes, and CD19^+^ B cells in PBMCs infected with E. coli MOI 1 based on negative control complementary antisense. **(D)** Summary plot of PBMCs populations positive for 16s rRNA, *n* = 3. (E) 16S rRNA positive PBMCs infected with *E. coli* MOI 1, analyzed by imagine flow.

**Fig S5: Detection of 16S rRNA in different populations of PBMCs in patients with CRC.**

Representative gating strategy of PBMCs for single cells, viability dye, surface staining for CD3, CD4, CD8, CD14, CD19 and 16S rRNA probe and complementary antisense probe.

**Fig S6: Positive cells for 16S rRNA in different populations of PBMCs in patients with CRC.**

**(A)** Gating strategy for CD4^+^ T cells, CD8^+^ T cells, CD14^+^ monocytes, and CD19^+^ B cells was performed based on antisense template control. As analyzed by imaging flow cytometry, CRC donor shows an increased number of positive cells. Graphs present the number of positive cells in each population. **(B)** Representative pictures of PBMCs from positive cells for 16S rRNA, and CD3, CD4, CD8, and CD14 cell markers.

**Fig S7: Positive cells for 16S rRNA in different populations of PBMCs in patients with IBD.**

**(A)** Gating strategy for CD4^+^ T cells, CD8^+^ T cells, CD14^+^ monocytes, and CD19^+^ B cells was performed based on antisense template control. IBD patient shows increased positive cells analyzed by imaging flow cytometry. Graphs presents several positive cells in each population. **(B)**Representative pictures of PBMCs from positive cells for 16S rRNA and CD3, CD4, CD8, and CD14 cell markers.

**Fig. S8: Statistically significant bacterial species in PBMCs and tissues.**

The heatmap presents the relative abundance of statistically significant species across PBMCs, their corresponding tissue, and their prevalence in the patient samples. Black squares represent significance in the corresponding comparison. Two-way ANOVA with disease and tissue type as factors followed by pairwise Wilcoxon tests with FDR adjustments for multiple testing and an adjusted p-value less than 0.05 is considered significant.

**Fig. S9: Validation of species-specific primers with pure bacterial DNA.**

Validation of primers for **(A)** *Anaerostipes hadrus*, **(B)** *Bifidobacterium adolescentis*, **(C)** *Collinsella aerofaciens* and **(D)** *Faecelibacterium prausnitzii* using 5 ng of pure bacterial DNA for each species. The left site shows the amplification plot while on the right the melt curve is illustrated.

**Fig. S10: Detection of species-specific DNA using qPCR.**

Standard curve for each primer, showing the Ct values and the respective log10 transformed DNA concentration from pure bacterial species ranging from 50 ng to 5 pg.

**Fig. S11: Altered metabolites in serum samples from patients with IBD or CRC compared to healthy controls.**

A heatmap displaying the top 156 altered metabolites, defined by an adjusted p-value of less than 0.0001, after an ANOVA statistical test between samples from patients with CRC (*n =* 28), IBD (*n* = 10), and healthy controls (*n =* 10).

**Fig. S12: Distinct pattern of bacterial translocation across disease types.**

In the intestinal tissue from patients with IBD and CRC, but not in their PBMCs, there is a high abundance of *Ruminococci*, *e*.*g*., the mucolytic *Ruminococcus torques*, or *Bacteroidaceae*, such as *Bacteroides fragilis* or *B. uniformis*. Those mucus degrading species might contribute to the aggravation of the intestinal barrier defects in such diseases. This will then allow additional bacteria to invade deeper into the intestinal mucosa and submucosa. Further, there is a high abundance of *Lachnospiraceae*, *e*.g*., Anaerostipes hadrus, Blautia wexlerae or Roseburia intestinalis* and *Faecalibacteria*, in the intestinal tissue as well as in the PBMCs of the patients. In the PBMCs from patients with a severe intestinal barrier defect, there are also high abundances of pathogenic bacteria, *e*.*g*., *Clostridioides difficile* or *Haemophilus parainfluenzae*. Some of the mentioned species were also detectable in the liver metastasis of the patients with CRC. Those observations might support the concept of translocation of bacterial remnants from the intestine into PBMCs of patients with CRC and IBD due to a defective intestinal epithelial barrier.

## Materials and Methods

### Patient Recruitment

We included patients with IBD and CRC. CRC patients diagnosed with Union for International Cancer Control (UICC) stages I-IV and a minimum of 18 years of age were included. Additionally, 20 healthy control individuals were included. Ethical approval for collecting and analyzing tissue and blood specimens from patients with CRC or IBD and healthy control individuals was received by the Cantonal Ethics Committee of the Canton Zürich (BASEC-No.: 2022-02036, BASEC-No.: 2019-02277, EK-1755/PB_2019-00169). Patients were recruited either from the University Hospital Zurich (USZ) or from the University Hospital Basel (USB). All participants provided written informed consent before sample collection. The control subjects were healthy individuals, e.g. undergoing CRC screening or blood donation at the Blood Donation Center, SRK Zurich, Switzerland. Control subjects were not formally age-matched. All samples from patients with CRC or IBD were taken after diagnosis, meaning that all patients had a confirmed diagnosis at the time of biosampling. Samples from patients with CRC were usually taken at time of cancer surgery. Samples from patients with IBD were taken during disease course. Once collected, the samples were treated identically between cases and controls with respect to timing, shipping, lab handling, processing.

### Tissue preparation

After surgical resection of human tissues, specimens were assessed by a pathologist. Tumor samples, inflamed and unaffected adjacent colon or liver samples were identified by the pathologist. Native tumor tissue and unaffected colon and liver tissue were temporally kept in sodium chloride (NaCl) until snap frozen in liquid nitrogen and stored at −80°C.

### Isolation of human PBMCs

Peripheral blood was collected in ethylenediaminetetraacetic acid (EDTA) (Beckton Dickinson (BD), 367525)-containing tubes and was processed within 1-2 h of blood draw. PBMCs were isolated using density gradient centrifugation with Pancoll-Paque at 1.077 g/mL (PanBiotech, PANP04-601000). Briefly, blood was diluted with 2% Fetal Calf Serum (FCS) (Biowest,, #S181T) in Phosphate-Buffered Saline (PBS) (Thermofisher, 10010056) and gently transferred on top of the density gradient medium layer in SepMate tubes (Stemcell, 85450). Samples were centrifuged at 1000 x *g* for 10 minutes at room temperature (RT), 22°C with the brake on 7. The resulting layer of mononuclear cells was transferred to a new tube, washed with 2% FCS in PBS, and centrifuged at 600 x *g* for 15 minutes with the brake on 7, at RT, followed by one additional wash with 2% FCS in PBS at 300 x *g* with the brake on 7 for 10 minutes. In FCS, cells were frozen in 10% dimethylsulfoxide (DMSO) (Sigma-Aldrich, 41640) and stored at - 80°C.

### Infection of human PBMCs with E. coli

The PBMCs were plated in 6-well plate (5 × 106 cells in 5 ml RPMI medium, Gibco) overnight at 37℃ in 5% CO2 atmosphere. Next, PBMCs were treated with antibiotics (gentamicin (100 μg/ml), doxycycline (5 μg/ml)) as a control or infected with different MOI (0, 0.5, 1, 2) of *E. coli* for 2 h. After infection, PBMCs were collected and washed three times with PBS and followed the 16S rRNA FISH staining.

### 16S rRNA FISH staining

The PBMCs were washed three times with PBS. Next, cells were stained for zombie violet fixable viability dye (Biolegend) 30 min at 4℃, and permeabilized for 20 min at 4℃ with intracellular staining permeabilization wash buffer and fixation buffer (Biolegend). Next, cells were washed with 2x saline-sodium citrate (SSC) buffer (20x SSC buffer, Roth, 1054.1) and stained with a 16Ss rRNA FISH probe EUB338 (5’-GCTGCCTCCCGTA-GGAGT-3), Cy5 (Microsynth) or an antisense probe (5’-ACTCCTACGG-GAGGCAGC-3’), Cy5 (Microsynth) in hybridization buffer (25% formamide Huberlab, A2156.0500), 10% dextran sulfate (ThermoFisher, 15415089), 0.5% bovine serum albumin (BSA) (Pan-Biotec, PANP06-1391500), 2x SSC buffer) and incubated overnight at 37℃ in 5% CO2 atmosphere. Next, cells were washed with 2x SSC buffer. After washing with MACS buffer (Miltenyi Biotec, 130091221) cells were stained with the surface markers CD3 (BV605), CD4 (PE), CD8 (PercpCy5), CD14 (APCCy7) and CD19 (PEDazzle) (Biolegend) for 30 min at RT. Flow cytometric acquisition was performed on the LSR II Fortessa-4L (BD) flow cytometer or Image stream X Mark II imaging flow cytometry (AMNIS). Flow cytometry data were analyzed using the FlowJo software (BD), Imaging flow cytometry data were analyzed using IDEAS software (Cytek Biosciences).

### Nucleic acid extraction from PBMCs and tissue samples

Nucleic acid extraction was performed on tissue samples, PBMCs, and the control buffer samples (serving as negative control). Respective sequencing reads are shown in Table S2. No human DNA sequence depletion or enrichment of microbial or viral DNA was performed. In detail, PBMCs and buffer samples were centrifuged for 10 min at 6000 x *g* and supernatant discarded. Microbial DNA from cell pellets and tissue samples was extracted with the ZymoBIOMICS DNA miniprep kit (Zymo Research, D4300), following the manufacturer’s instructions. Instead of Zymo-Spin IICR single columns, an E-Z 96 DNA plate (Omega Bio-Tek IBD96-02) was used, and centrifugation speed was reduced to 6000 x *g*. DNA was eluted in 60 µL elution buffer and quantified using the PicoGreen dsDNA Assay (Thermo Fisher Scientific) or Pico488 dsDNA Assay (Lumiprobe).

### Shotgun Metagenomics Sequencing

Shotgun metagenomics sequencing was performed at Microsynth AG, Balgach, CH. The library preparation utilized the Illumina TruSeq library method with unique dual indexing (UDI). Sequencing was conducted on the Illumina NovaSeq platform, specifically on the SP4 chemistry, using a paired-end 2×100 sequencing strategy. The library preparation workflow included a comprehensive quality control (QC) step for the samples, library preparation, and a final library QC assessment. Libraries were quantified and equimolarly pooled to ensure uniform representation. The Illumina NovaSeq S4 flow cell used for DNA sequencing had a specified throughput of 16,000 million passed filter reads with an average of 30 million reads per sample. We analyzed the collected metagenomics dataset using the same settings. Raw reads were trimmed for quality using fastp^52^ and retained high-quality sequences, and then human host reads were subtracted by mapping the reads with the human reference genome (GRCh38) using Bowtie2^53^. Fastq files were assessed for quality control using the FASTQC application. Samples were subjected to sequencing across two distinct batches. To account for potential batch effects that could confound the analysis, we employed the ConQuR method40, a robust statistical technique based on conditional quantile regression designed specifically for the normalization of microbiome data.

### Taxonomic Classification

Taxonomic classification was performed using the Kraken2 pipeline (version 2.0.8)^54^. Clean reads were classified taxonomically against the PlusPF Kraken2 database using the Kraken2 classification tool with default parameters. All human reads were excluded and the relative abundance was calculated based on the successful alignment against the bacteria db. Functional profiling was performed using HUMAnN3^55^, by mapping the sequences against the UniRef90 database after lowering the threshold of MetaPhlan’s 3 intermediary taxonomy level to 0.0001. Since HUMAnN3 does not natively support paired-end reads as input, the corresponding FASTQ files were run in separate runs, and the average of matched results between forward and reverse reads were considered for further analysis.

### Filtering and cleaning the data

To effectively decontaminate 7144 bacterial taxa detected in our microbiome study, especially because we involve low-biomass samples, a multi-faceted approach combining ecological plausibility, statistical analyses, and stringent experimental controls is essential. Samples were collected and processed using sterile techniques in a controlled environment, with multiple negative controls included, including water, buffer, and freezing medium **(Table S2)** to detect potential contamination at each stage. DNA was extracted in two different batches using different kit lots to identify batch effects. To ensure ecological plausibility, we considered only the species that were detected by 4 fold (10 reads) of average bacterial reads (2.25) in blank samples. Moreover, the species were only considered for further filtering if those were detected with more than ten reads in at least 3 patients in one of the diseases/tissues/PBMCs, reducing the species number to 738. Then we performed hierarchical correlation clustering to detect clusters with possible contamination **(Fig. S1A-B)**. Principal component analysis was used to identify and visualize batch effects. Finally, to achieve genuine signals we removed cited reagent contamination to end up with 208 species detected in biopsies and PBMCs^24,25^.

### Species specific qPCR

Quantitative real-time polymerase chain reaction (qPCR) amplification was performed using species specific primers for *Anaerostipes hadrus* (5’-CTTTAGTAGCCAGCATATAAGG -3’, 5’-TTGCTCACTCTCACGAGGCT-3’)^56^, *Bifidobacterium adolescentis* (5’-CTCCAGTTGGATGCATGTC-3’, 5’-CGAAGGCTTGCTCCCAGT-3’)^57^, *Collinsella aerofaciens* (5’-CCCGACGGGAGGGGAT-3, 5’-CTTCTGCAGGTACAGTCTTGAC-3’)^58^, and *Faecalibacterium prausnitzii* (5’-GGAGGAAGAAGGTCTTCGG-3’, 5’-AATTCCGCCTACCTCTGCACT-3’)^59^. Pure bacterial DNA (DSM 23942, DSM 20083, DSM 3979, DSM107838) was used to perform standard curve analysis **(Fig. S9)** and to validate primers and **(Fig. S10A-D)**. One qPCR reaction in a total volume of 10 uL consisted of 5 uL of Power SYBR Green PCR Mastermix (Thermo Fisher Scientific, 4367659), 0.1 uL of each primer at 100 uM, nuclease free water (Promega, P119C) and the respective DNA sample. 100 ng of whole DNA (including eukaryotic and possible microbial DNA) isolated from PBMCs, adjacent CRC and CRC as well as non-inflamed and inflamed IBD tissue was used as input per reaction. We used remaining isolated DNA from the same patient samples on which previously metagenomic sequencing was performed. qPCR was run in triplicates and non-template controls (NTCs), which included the same reagents but used water in place of DNA input, were incorporated into each qPCR run. qPCR was run on Quant Studio 6 Flex (278861872) and thermal cycler settings were adapted according to the manufacturer’s instructions and an annealing temperature of 68°C.

### Metabolomics

Blood was collected in serum vacutainers (BD, 367896), inverted 5 times, left at least 30 min at RT for clotting and then centrifuged at 2000 x *g* for 10 minutes at RT. Subsequently obtained serum was snap frozen and stored at −80°C until further processing. Metabolomic analysis was carried out with a high-resolution mass spectrometer (Agilent QTOF 6550) described previously by Fuhrer *et al.*^60^, and we used MetaboAnalystR package version 3.0.3 for the statistical analysis^61^. The normalized intensities by the median for each sample were mean-centered and divided by the standard deviation of each variable of all annotated features. Significance analysis was done by ANOVA followed by Tukey-HSD post-hoc analysis. Principle component analysis (PCA) was used to visualize sample variance. Heatmap visually presented the hierarchical clustering using the Euclidean method for distance calculation and Ward’s linkage for clustering. Metabolite set enrichment analysis (MSEA) was carried out against the “main” data set, and a generalized linear model was used during the quantitative enrichment analysis. The correlation between the significant metabolites and enriched pathways was conducted using the corrr package in R and the sanky plot was generated using (networkD3) package.

### Statistical analysis

Statistical analyses were performed in R (version 4.4.3). We employed two methods for normalization based on the type of analysis. We applied rarefaction to subsample down to 20 reads per sample for diversity analyses. Rarefaction was employed to standardize sequencing depth among samples, preventing diversity comparisons from being skewed by varying sampling efforts or sequencing depths. By establishing a minimum sequencing depth of 20 reads per sample, we ensure that only those with adequate data are considered. Given that we are identifying very low biomass, we lowered the depth to balance the retention of as many samples as possible while maintaining reliable data for thorough diversity analysis. Consequently, several samples were excluded from the analysis due to their sequencing depths not meeting the set threshold. Alpha-diversity (the number of observed genera and Chao1 indices (richness), and beta-diversity (Bray-Curtis and Jaccard dissimilarities) indices were generated using vegdist and diversity functions in the vegan R package (version 2.3-2). Principal coordinate analysis (PCoA) was performed using the phyloseq package based on the Bray-Curtis. Permutational variance analysis (PERMANOVA) was conducted using the adonis2 function from R’s vegan package. Differential abundance was assessed using relative abundance. This approach accounts for the differences in total read counts between PBMC and tissue samples, enabling meaningful comparisons of bacterial composition. Either the Wilcoxon test for 2 group comparison was used or the Kruskal-Wallis test followed by Wilcoxon test for pairwise comparisons and p value adjusted using FDR for family error correction. Linear discriminant analysis effect size (LEFSe)^62^ was used to identify differentially abundant bacteria species between two or more groups, and the linear discriminant analysis (LDA) score was obtained. The correlation between the significant metabolites and enriched pathways was conducted using the corrr package in R and the sanky plot was generated using (networkD3) package.

## Main References

1 Zhang, Y. et al. Fusobacterium nucleatum promotes colorectal cancer cells adhesion to endothelial cells and facilitates extravasation and metastasis by inducing ALPK1/NF-kappaB/ICAM1 axis. Gut Microbes 14, 2038852 (2022). 10.1080/19490976.2022.2038852

2 Pleguezuelos-Manzano, C. et al. Mutational signature in colorectal cancer caused by genotoxic pks(+) E. coli. Nature 580, 269–273 (2020). 10.1038/s41586-020-2080-8

3 Galeano Nino, J. L., et al. Effect of the intratumoral microbiota on spatial and cellular heterogeneity in cancer. Nature 611, 810–817 (2022). 10.1038/s41586-022-05435-0

4 Zheng, D., Liwinski, T. & Elinav, E. Interaction between microbiota and immunity in health and disease. Cell Res 30, 492–506 (2020). 10.1038/s41422-020-0332-7

5 Anderson, N. M. & Simon, M. C. The tumor microenvironment. Curr Biol 30, R921–R925 (2020). 10.1016/j.cub.2020.06.081

6 Shan, Y., Lee, M. & Chang, E. B. The Gut Microbiome and Inflammatory Bowel Diseases. Annu Rev Med 73, 455–468 (2022). 10.1146/annurev-med-042320-021020

7 Bullman, S. et al. Analysis of Fusobacterium persistence and antibiotic response in colorectal cancer. Science 358, 1443–1448 (2017). 10.1126/science.aal5240

8 Tan, C. C. S. et al. No evidence for a common blood microbiome based on a population study of 9,770 healthy humans. Nat Microbiol 8, 973–985 (2023). 10.1038/s41564-023-01350-w

9 Hou, K. et al. Microbiota in health and diseases. Signal Transduct Target Ther 7, 135 (2022). 10.1038/s41392-022-00974-4

10 Xu, X. et al. The gut metagenomics and metabolomics signature in patients with inflammatory bowel disease. Gut Pathog 14, 26 (2022). 10.1186/s13099-022-00499-9

11 Proctor, L. M. et al. The Integrative Human Microbiome Project. Nature 569, 641–648 (2019). 10.1038/s41586-019-1238-8

12 Huttenhower, C. et al. Structure, function and diversity of the healthy human microbiome. Nature 486, 207–214 (2012). 10.1038/nature11234

13 Methé, B. A. et al. A framework for human microbiome research. Nature 486, 215–221 (2012). 10.1038/nature11209

14 Forbes, J. D., Van Domselaar, G. & Bernstein, C. N. Microbiome Survey of the Inflamed and Noninflamed Gut at Different Compartments Within the Gastrointestinal Tract of Inflammatory Bowel Disease Patients. Inflamm Bowel Dis 22, 817–825 (2016). 10.1097/MIB.0000000000000684

15 Darfeuille-Michaud, A. et al. High prevalence of adherent-invasive Escherichia coli associated with ileal mucosa in Crohn’s disease. Gastroenterology 127, 412–421 (2004). 10.1053/j.gastro.2004.04.061

16 Read, E., Curtis, M. A. & Neves, J. F. The role of oral bacteria in inflammatory bowel disease. Nat Rev Gastroenterol Hepatol 18, 731–742 (2021). 10.1038/s41575-021-00488-4

17 Montalban-Arques, A. et al. Commensal Clostridiales strains mediate effective anti-cancer immune response against solid tumors. Cell Host Microbe 29, 1573–1588 e1577 (2021). 10.1016/j.chom.2021.08.001

18 Kostic, A. D. et al. Fusobacterium nucleatum potentiates intestinal tumorigenesis and modulates the tumor-immune microenvironment. Cell Host Microbe 14, 207–215 (2013). 10.1016/j.chom.2013.07.007

19 Wirbel, J. et al. Meta-analysis of fecal metagenomes reveals global microbial signatures that are specific for colorectal cancer. Nat Med 25, 679–689 (2019). 10.1038/s41591-019-0406-6

20 Thomas, A. M. et al. Metagenomic analysis of colorectal cancer datasets identifies cross-cohort microbial diagnostic signatures and a link with choline degradation. Nat Med 25, 667–678 (2019). 10.1038/s41591-019-0405-7

21 Dejea, C. M. et al. Patients with familial adenomatous polyposis harbor colonic biofilms containing tumorigenic bacteria. Science 359, 592–597 (2018). 10.1126/science.aah3648

22 Bertocchi, A. et al. Gut vascular barrier impairment leads to intestinal bacteria dissemination and colorectal cancer metastasis to liver. Cancer Cell 39, 708–724 e711 (2021). 10.1016/j.ccell.2021.03.004

23 Messaritakis, I. et al. The Prognostic Value of the Detection of Microbial Translocation in the Blood of Colorectal Cancer Patients. Cancers (Basel) 12 (2020). 10.3390/cancers12041058

24 de Goffau, M. C. et al. Recognizing the reagent microbiome. Nat Microbiol 3, 851–853 (2018). 10.1038/s41564-018-0202-y

25 Kennedy, K. M. et al. Questioning the fetal microbiome illustrates pitfalls of low-biomass microbial studies. Nature 613, 639–649 (2023). 10.1038/s41586-022-05546-8

26 Kitamura, T., Qian, B. Z. & Pollard, J. W. Immune cell promotion of metastasis. Nat Rev Immunol 15, 73–86 (2015). 10.1038/nri3789

27 Zhai, W., Wu, F., Zhang, Y., Fu, Y. & Liu, Z. The Immune Escape Mechanisms of Mycobacterium Tuberculosis. Int J Mol Sci 20 (2019). 10.3390/ijms20020340

28 Eisele, N. A. et al. Salmonella require the fatty acid regulator PPARdelta for the establishment of a metabolic environment essential for long-term persistence. Cell Host Microbe 14, 171–182 (2013). 10.1016/j.chom.2013.07.010

29 Amar, J. et al. Intestinal mucosal adherence and translocation of commensal bacteria at the early onset of type 2 diabetes: molecular mechanisms and probiotic treatment. EMBO Mol Med 3, 559–572 (2011). 10.1002/emmm.201100159

30 Abed, J. et al. Colon Cancer-Associated Fusobacterium nucleatum May Originate From the Oral Cavity and Reach Colon Tumors via the Circulatory System. Front Cell Infect Microbiol 10, 400 (2020). 10.3389/fcimb.2020.00400

31 Gihawi, A. et al. Major data analysis errors invalidate cancer microbiome findings. mBio 14, e0160723 (2023). 10.1128/mbio.01607-23

32 Geller, L. T. et al. Potential role of intratumor bacteria in mediating tumor resistance to the chemotherapeutic drug gemcitabine. Science 357, 1156–1160 (2017). 10.1126/science.aah5043

33 Amann, R. I. et al. Combination of 16S rRNA-targeted oligonucleotide probes with flow cytometry for analyzing mixed microbial populations. Appl Environ Microbiol 56, 1919–1925 (1990). 10.1128/aem.56.6.1919-1925.1990

34 Markman, J. L. & Shiao, S. L. Impact of the immune system and immunotherapy in colorectal cancer. J Gastrointest Oncol 6, 208–223 (2015). 10.3978/j.issn.2078-6891.2014.077

35 Ai, D. et al. Identifying Gut Microbiota Associated With Colorectal Cancer Using a Zero-Inflated Lognormal Model. Front Microbiol 10, 826 (2019). 10.3389/fmicb.2019.00826

36 Salazar, A. M., Aparicio, R., Clark, R. I., Rera, M. & Walker, D. W. Intestinal barrier dysfunction: an evolutionarily conserved hallmark of aging. Dis Model Mech 16 (2023). 10.1242/dmm.049969

37 Ghosh, T. S., Shanahan, F. & O’Toole, P. W. The gut microbiome as a modulator of healthy ageing. Nat Rev Gastroenterol Hepatol 19, 565–584 (2022). 10.1038/s41575-022-00605-x

38 Lloyd-Price, J. et al. Multi-omics of the gut microbial ecosystem in inflammatory bowel diseases. Nature 569, 655–662 (2019). 10.1038/s41586-019-1237-9

39 Human Microbiome Project, C. Structure, function and diversity of the healthy human microbiome. Nature 486, 207–214 (2012). 10.1038/nature11234

40 Kim, M. Y. et al. CD4(+)CD3(−) accessory cells costimulate primed CD4 T cells through OX40 and CD30 at sites where T cells collaborate with B cells. Immunity 18, 643–654 (2003). 10.1016/s1074-7613(03)00110-9

41 Parel, Y. & Chizzolini, C. CD4+ CD8+ double positive (DP) T cells in health and disease. Autoimmunity reviews 3, 215–220 (2004). 10.1016/j.autrev.2003.09.001

42 Desfrancois, J. et al. Double positive CD4CD8 alphabeta T cells: a new tumor-reactive population in human melanomas. PLoS One 5, e8437 (2010). 10.1371/journal.pone.0008437

43 Quandt, D., Rothe, K., Scholz, R., Baerwald, C. W. & Wagner, U. Peripheral CD4CD8 double positive T cells with a distinct helper cytokine profile are increased in rheumatoid arthritis. PLoS One 9, e93293 (2014). 10.1371/journal.pone.0093293

44 Baba, T. et al. CD4+/CD8+ macrophages infiltrating at inflammatory sites: a population of monocytes/macrophages with a cytotoxic phenotype. Blood 107, 2004–2012 (2006). 10.1182/blood-2005-06-2345

45 Pallett, L. J. et al. Tissue CD14(+)CD8(+) T cells reprogrammed by myeloid cells and modulated by LPS. Nature 614, 334–342 (2023). 10.1038/s41586-022-05645-6

46 Mutignani, M. et al. Blood Bacterial DNA Load and Profiling Differ in Colorectal Cancer Patients Compared to Tumor-Free Controls. Cancers (Basel) 13 (2021). 10.3390/cancers13246363

47 Liu, D. et al. Anaerostipes hadrus, a butyrate-producing bacterium capable of metabolizing 5-fluorouracil. mSphere 9, e0081623 (2024). 10.1128/msphere.00816-23

48 Vujkovic-Cvijin, I. et al. The systemic anti-microbiota IgG repertoire can identify gut bacteria that translocate across gut barrier surfaces. Sci Transl Med 14, eabl3927 (2022). 10.1126/scitranslmed.abl3927

49 Genua, F., Raghunathan, V., Jenab, M., Gallagher, W. M. & Hughes, D. J. The Role of Gut Barrier Dysfunction and Microbiome Dysbiosis in Colorectal Cancer Development. Front Oncol 11, 626349 (2021). 10.3389/fonc.2021.626349

50 Choi, Y. et al. Immune checkpoint blockade induces gut microbiota translocation that augments extraintestinal antitumor immunity. Sci Immunol 8, eabo2003 (2023). 10.1126/sciimmunol.abo2003

51 Turner, J. R. Intestinal mucosal barrier function in health and disease. Nat Rev Immunol 9, 799–809 (2009). 10.1038/nri2653

## Method References

52 Chen, S., Zhou, Y., Chen, Y. & Gu, J. fastp: an ultra-fast all-in-one FASTQ preprocessor. Bioinformatics 34, i884–i890 (2018). 10.1093/bioinformatics/bty560

53 Langmead, B., Wilks, C., Antonescu, V. & Charles, R. Scaling read aligners to hundreds of threads on general-purpose processors. Bioinformatics 35, 421–432 (2019). 10.1093/bioinformatics/bty648

54 Wood, D. E., Lu, J. & Langmead, B. Improved metagenomic analysis with Kraken 2. Genome Biol 20, 257 (2019). 10.1186/s13059-019-1891-0

55 Beghini, F. et al. Integrating taxonomic, functional, and strain-level profiling of diverse microbial communities with bioBakery 3. Elife 10 (2021). 10.7554/eLife.65088

56 Louis, P., Young, P., Holtrop, G. & Flint, H. J. Diversity of human colonic butyrate-producing bacteria revealed by analysis of the butyryl-CoA:acetate CoA-transferase gene. Environ Microbiol 12, 304–314 (2010). 10.1111/j.1462-2920.2009.02066.x

57 Matsuki, T., Watanabe, K., Tanaka, R. & Oyaizu, H. Rapid identification of human intestinal bifidobacteria by 16S rRNA-targeted species- and group-specific primers. FEMS Microbiol Lett 167, 113–121 (1998). 10.1111/j.1574-6968.1998.tb13216.x

58 Kassinen, A. et al. The fecal microbiota of irritable bowel syndrome patients differs significantly from that of healthy subjects. Gastroenterology 133, 24–33 (2007). 10.1053/j.gastro.2007.04.005

59 Ramirez-Farias, C. et al. Effect of inulin on the human gut microbiota: stimulation of Bifidobacterium adolescentis and Faecalibacterium prausnitzii. Br J Nutr 101, 541–550 (2009). 10.1017/S0007114508019880

60 Fuhrer, T., Heer, D., Begemann, B. & Zamboni, N. High-throughput, accurate mass metabolome profiling of cellular extracts by flow injection-time-of-flight mass spectrometry. Anal Chem 83, 7074–7080 (2011). 10.1021/ac201267k

61 Pang, Z., Chong, J., Li, S. & Xia, J. MetaboAnalystR 3.0: Toward an Optimized Workflow for Global Metabolomics. Metabolites 10 (2020). 10.3390/metabo10050186

62 Segata, N. et al. Metagenomic biomarker discovery and explanation. Genome Biol 12, R60 (2011). 10.1186/gb-2011-12-6-r60

