## Supplemental Figures S1-S12 for "Blood-borne immune cells carry low biomass DNA remnants of microbes in patients with colorectal cancer or inflammatory bowel disease"

A

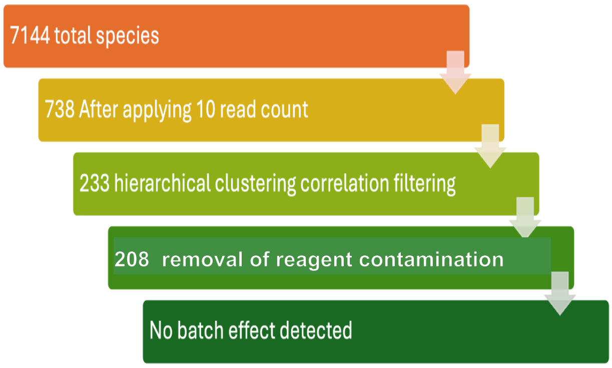
7144 total species
738 After applying 10 read count
233 hierarchical clustering correlation filtering
208 removal of reagent contamination
No batch effect detected

C

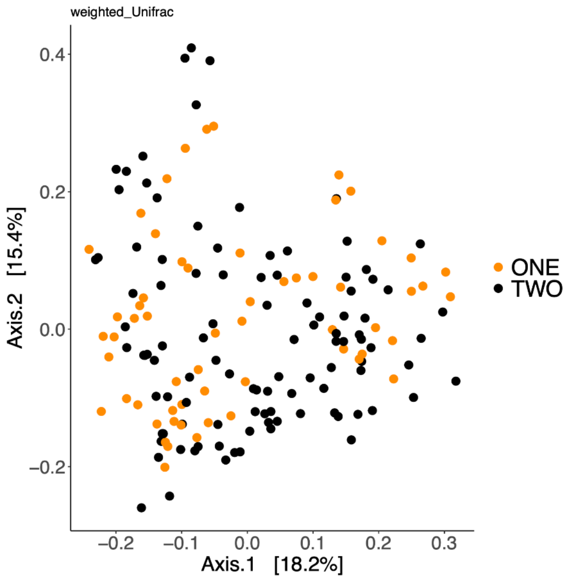
weighted_Unifrac
Axis.2 [15.4%]
Axis.1 [18.2%]
ONE
TWO
0.4
0.2
0.0
−0.2
−0.1
0.1
0.3

B

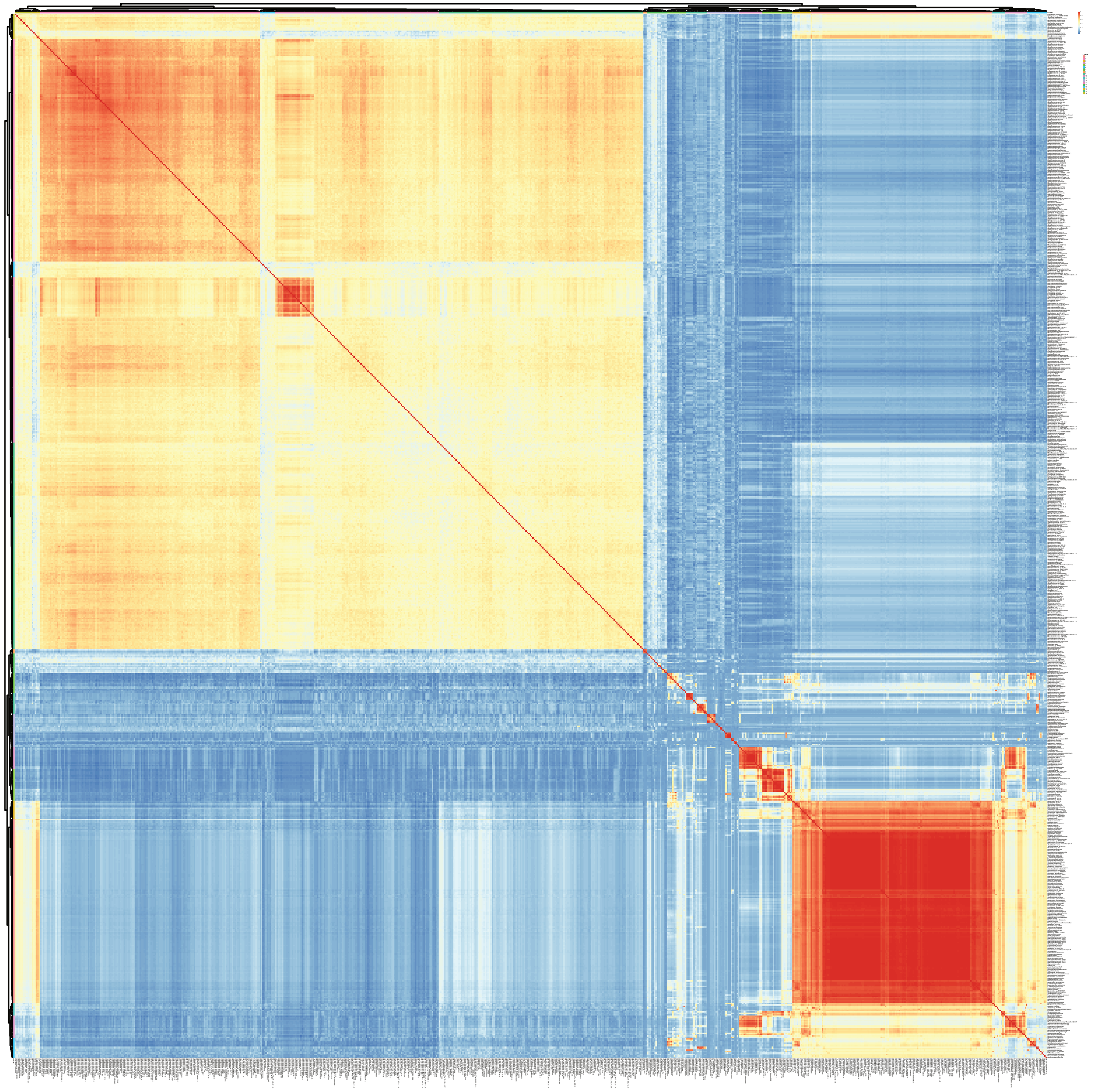

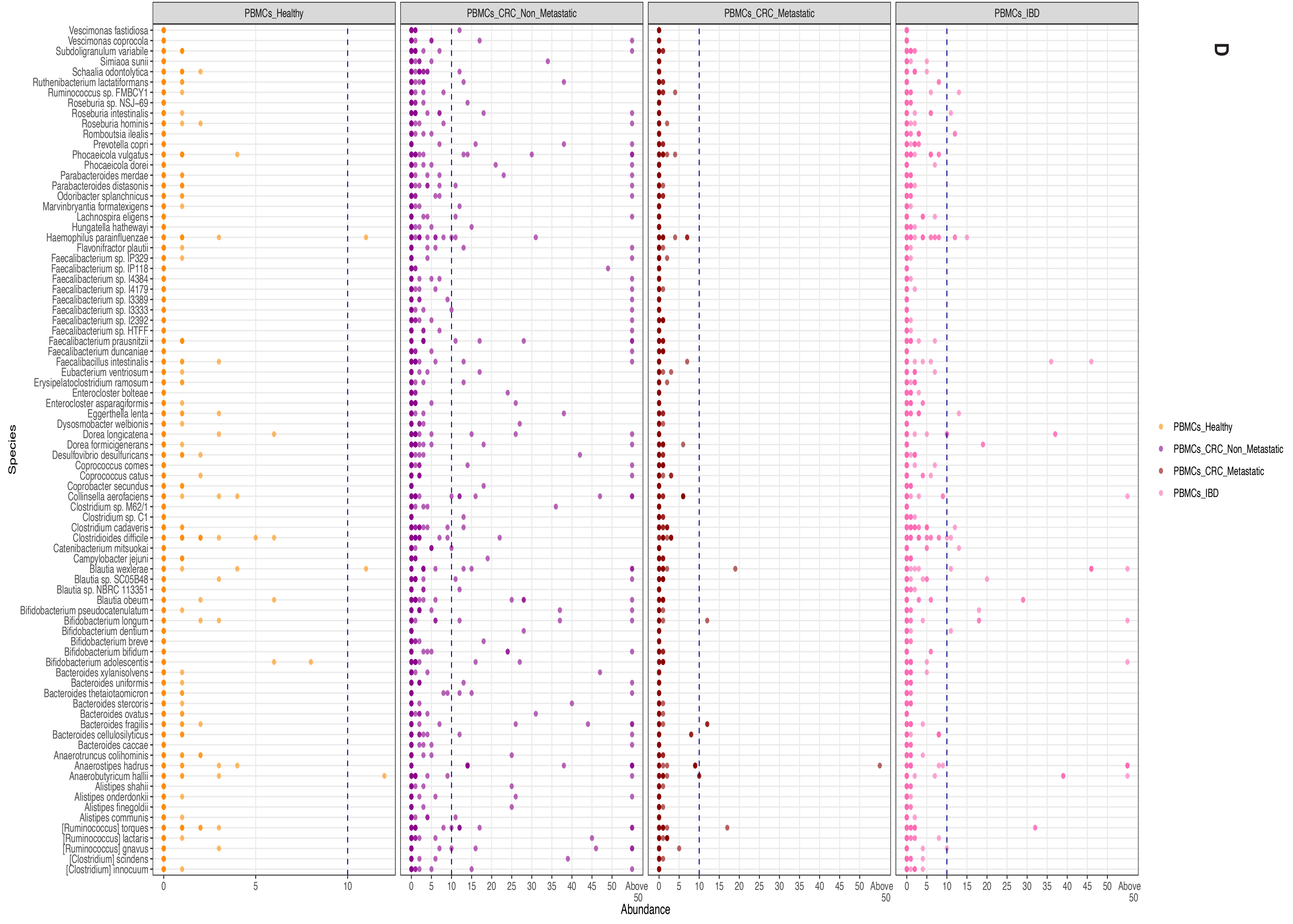
D
PBMCs_Healthy
PBMCs_CRC_Non_Metastatic
PBMCs_CRC_Metastatic
PBMCs_IBD
Species
Vescimonas fastidiosa
Vescimonas coprocola
Subdoligranulum variabile
Simiaoa sunii
Schaalia odontolytica
Ruthenibacterium lactatiformans
Ruminococcus sp. FMBCY1
Roseburia sp. NSJ-69
Roseburia intestinalis
Roseburia hominis
Romboutsia ilealis
Prevotella copri
Phocaeicola vulgatus
Phocaeicola dorei
Parabacteroides merdae
Parabacteroides distasonis
Odoribacter splanchnicus
Marvinbryantia formatexigens
Lachnospira eligens
Hungatella hathewayi
Haemophilus parainfluenzae
Flavonifractor plautii
Faecalibacterium sp. IP329
Faecalibacterium sp. IP118
Faecalibacterium sp. I4384
Faecalibacterium sp. I4179
Faecalibacterium sp. I3389
Faecalibacterium sp. I3333
Faecalibacterium sp. I2392
Faecalibacterium sp. HTFF
Faecalibacterium prausnitzii
Faecalibacterium duncaniae
Faecalibacillus intestinalis
Eubacterium ventriosum
Erysipelatoclostridium ramosum
Enterocloster bolteae
Enterocloster asparagiformis
Eggerthella lenta
Dysosmobacter welbionis
Dorea longicatena
Dorea formicigenerans
Desulfovibrio desulfuricans
Coprococcus comes
Coprococcus catus
Coprobacter secundus
Collinsella aerofaciens
Clostridium sp. M62/1
Clostridium sp. C1
Clostridium cadaveris
Clostridioides difficile
Catenibacterium mitsuokai
Campylobacter jejuni
Blautia wexlerae
Blautia sp. SC05B48
Blautia sp. NBRC 113351
Blautia obeum
Bifidobacterium pseudocatenulatum
Bifidobacterium longum
Bifidobacterium dentium
Bifidobacterium breve
Bifidobacterium bifidum
Bifidobacterium adolescentis
Bacteroides xylanisolvens
Bacteroides uniformis
Bacteroides thetaiotaomicron
Bacteroides stercoris
Bacteroides ovatus
Bacteroides fragilis
Bacteroides cellulosilyticus
Bacteroides caccae
Anaerotruncus colihominis
Anaerostipes hadrus
Anaerobutyricum hallii
Alistipes shahii
Alistipes onderdonkii
Alistipes finegoldii
Alistipes communis
[Ruminococcus] torques
[Ruminococcus] lactaris
[Ruminococcus] gnavus
[Clostridium] scindens
[Clostridium] innocuum
0
5
10
0
5
10
15
20
25
30
35
40
45
50
Above 50
Abundance
PBMCs_Healthy
PBMCs_CRC_Non_Metastatic
PBMCs_CRC_Metastatic
PBMCs_IBD

# S2

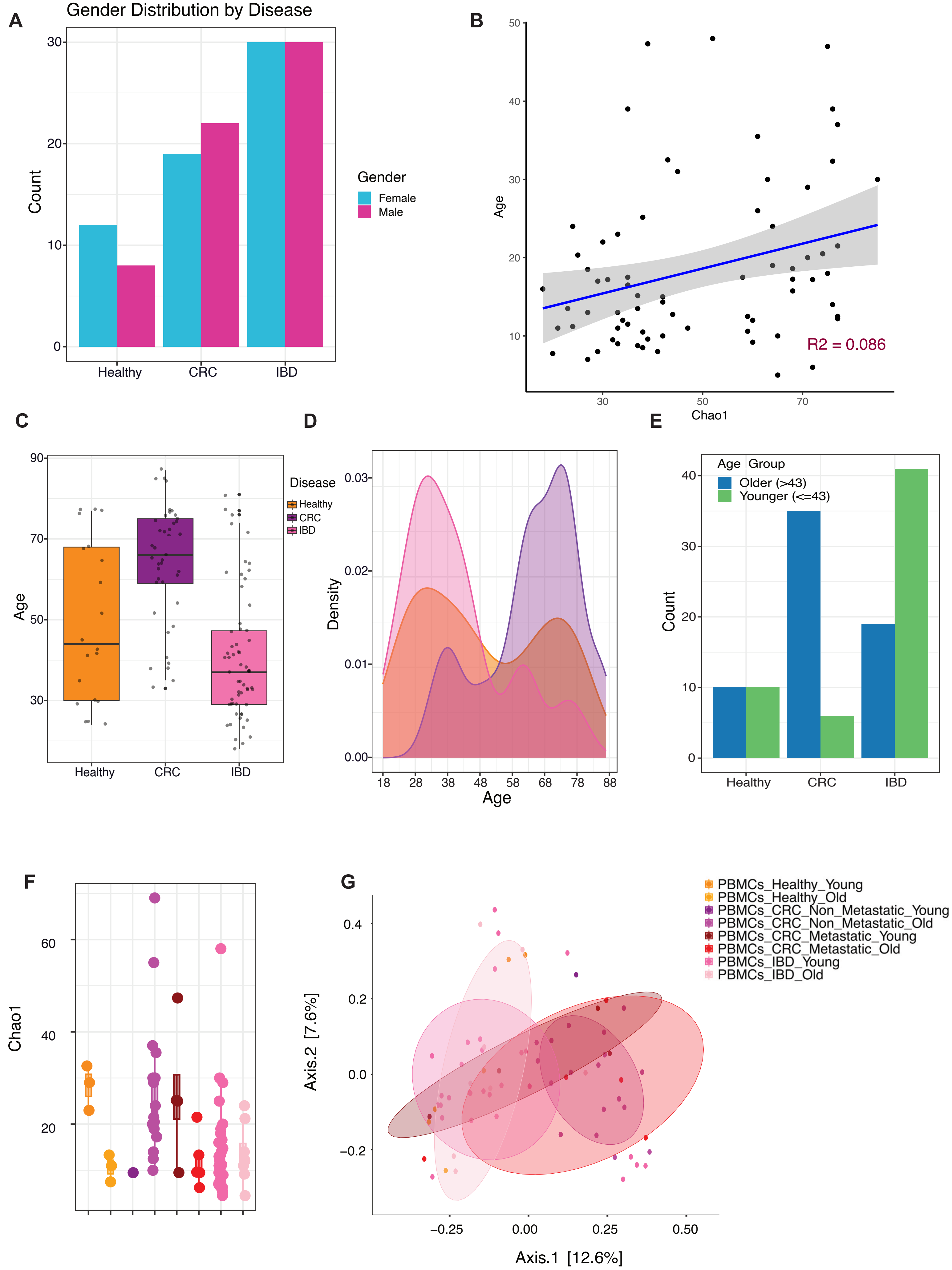
A
Gender Distribution by Disease
Count
Healthy
CRC
IBD
Gender
Female
Male
B
Age
Chao1
R2 = 0.086
C
Age
Disease
Healthy
CRC
IBD
D
Density
Age
E
Age_Group
Older (>43)
Younger (<=43)
Count
Healthy
CRC
IBD
F
Chao1
G
PBMCs_Healthy_Young
PBMCs_Healthy_Old
PBMCs_CRC_Non_Metastatic_Young
PBMCs_CRC_Non_Metastatic_Old
PBMCs_CRC_Metastatic_Young
PBMCs_CRC_Metastatic_Old
PBMCs_IBD_Young
PBMCs_IBD_Old
Axis.2 [7.6%]
Axis.1 [12.6%]

# S3

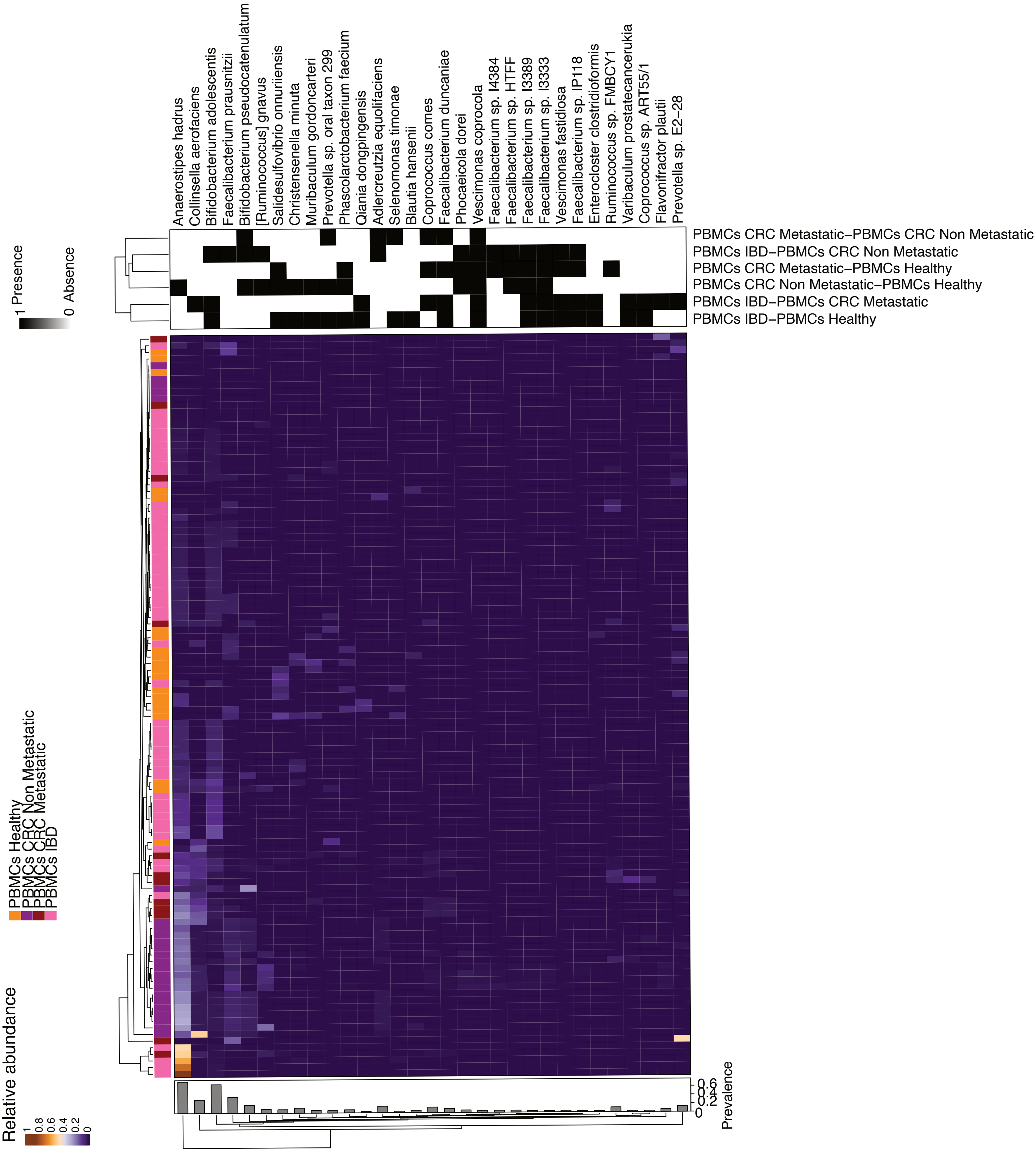
Anaerostipes hadrus
Collinsella aerofaciens
Bifidobacterium adolescentis
Faecalibacterium prausnitzii
Bifidobacterium pseudocatenulatum
[Ruminococcus] gnavus
Salidesulfovibrio onnuriiensis
Christensenella minuta
Muribaculum gordoncarteri
Prevotella sp. oral taxon 299
Phascolarctobacterium faecium
Qiania dongpingensis
Adlercreutzia equolifaciens
Selenomonas timonae
Blautia hansenii
Coprococcus comes
Faecalibacterium duncaniae
Phocaeicola dorei
Vescimonas coprocola
Faecalibacterium sp. I4384
Faecalibacterium sp. HTFF
Faecalibacterium sp. I3389
Faecalibacterium sp. I3333
Vescimonas fastidiosa
Faecalibacterium sp. IP118
Enterocloster clostridioformis
Ruminococcus sp. FMBCY1
Varibaculum prostatecancerukia
Coprococcus sp. ART55/1
Flavonifractor plautii
Prevotella sp. E2−28
PBMCs CRC Metastatic−PBMCs CRC Non Metastatic
PBMCs IBD−PBMCs CRC Non Metastatic
PBMCs CRC Metastatic−PBMCs Healthy
PBMCs CRC Non Metastatic−PBMCs Healthy
PBMCs IBD−PBMCs CRC Metastatic
PBMCs IBD−PBMCs Healthy
1 Presence
0 Absence
PBMCs Healthy
PBMCs CRC Non Metastatic
PBMCs CRC Metastatic
PBMCs IBD
Relative abundance
1
0.8
0.6
0.4
0.2
0
Prevalence
0.6
0.4
0.2
0

S4

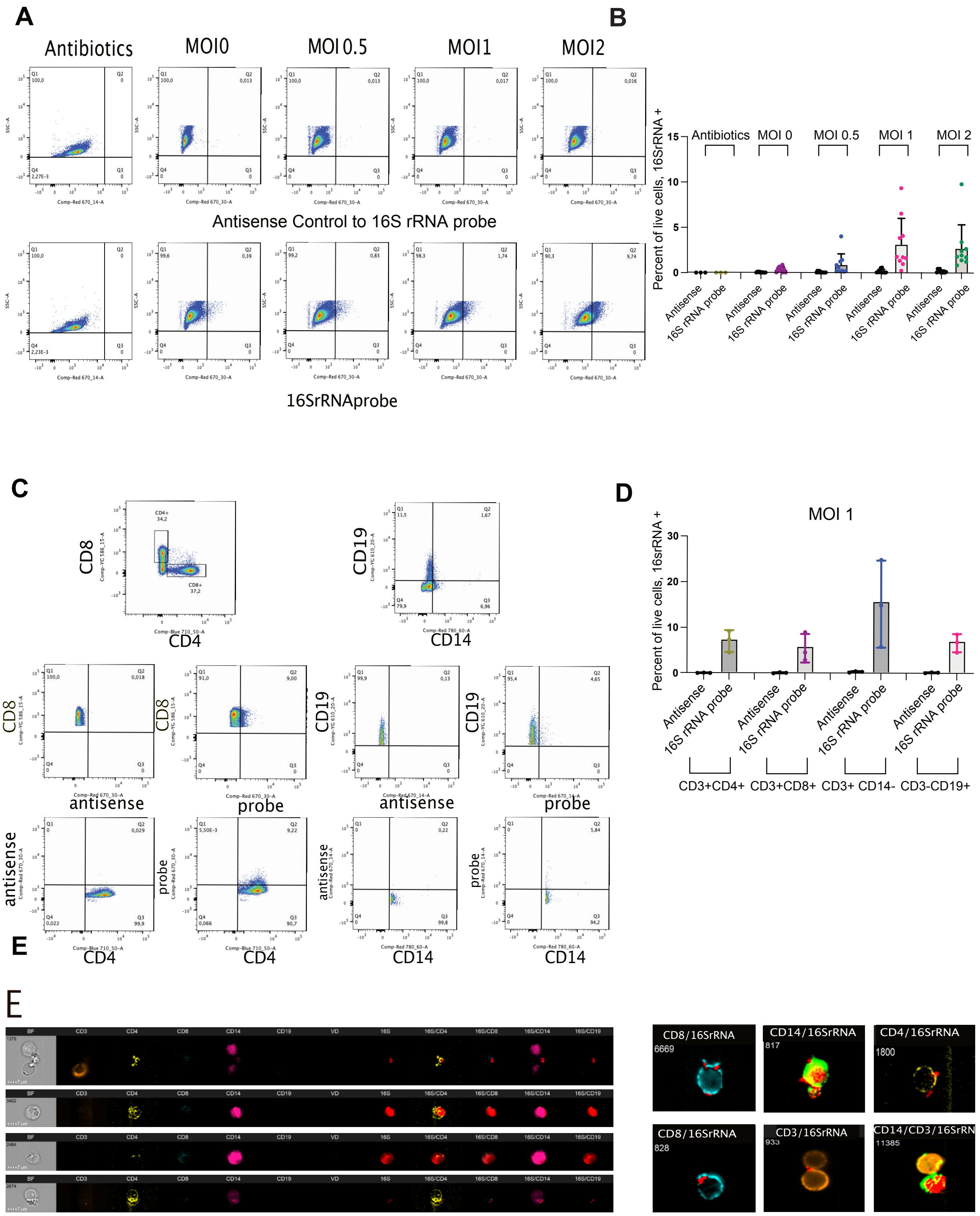
A
B
Antibiotics
MOI0
MOI 0.5
MOI1
MOI2
Antisense Control to 16S rRNA probe
16SrRNAprobe
Percent of live cells, 16SrRNA +
Antibiotics
MOI 0
MOI 0.5
MOI 1
MOI 2
Antisense
16S rRNA probe
C
D
MOI 1
CD8
CD4
CD19
CD14
antisense
probe
Percent of live cells, 16srRNA +
CD3+CD4+
CD3+CD8+
CD3+ CD14-
CD3-CD19+
E
CD8/16SrRNA
CD14/16SrRNA
CD4/16SrRNA
CD3/16SrRNA
CD14/CD3/16SrRN

**S5**

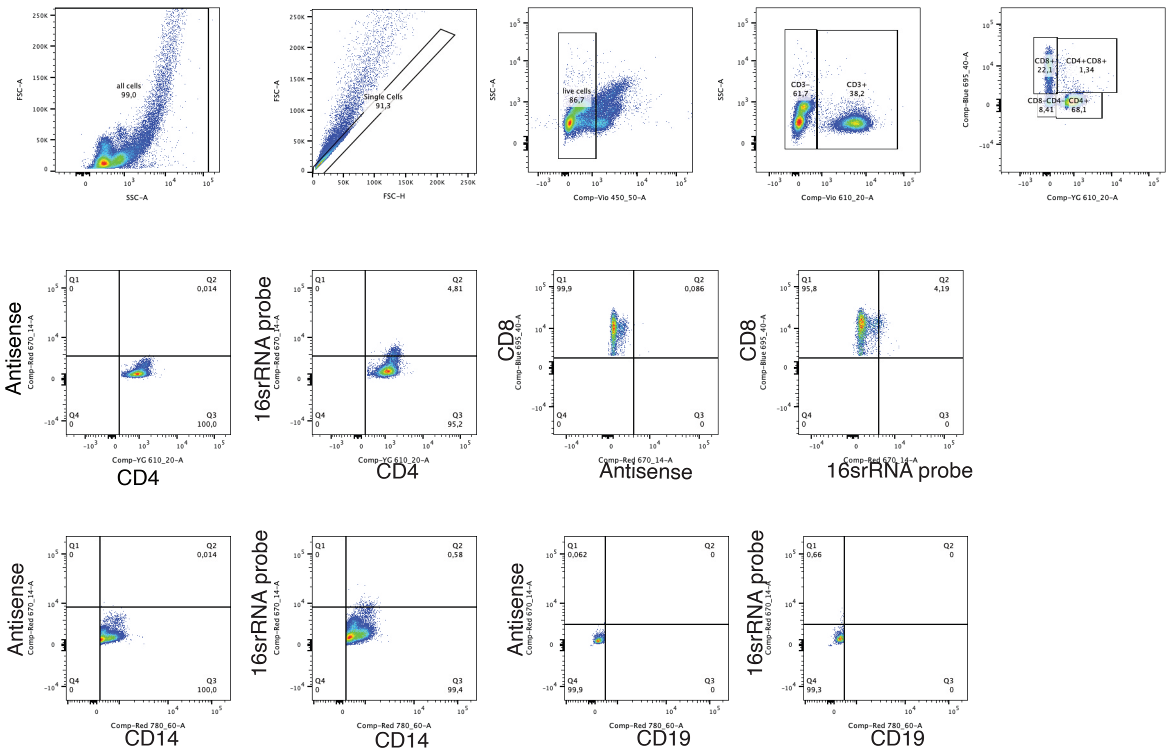

all cells
99,0
SSC-A
FSC-A
Single Cells
91,3
FSC-H
live cells
86,7
Comp-Vio 450_50-A
CD3-
61,7
CD3+
38,2
Comp-Vio 610_20-A
CD8+
22,1
CD4+CD8+
1,34
CD8-CD4-
8,41
CD4+
68,1
Comp-YG 610_20-A
Comp-Blue 695_40-A
Antisense
16srRNA probe
CD4
CD8
CD14
CD19
Comp-Red 670_14-A
Comp-Red 780_60-A

# S6

**A**

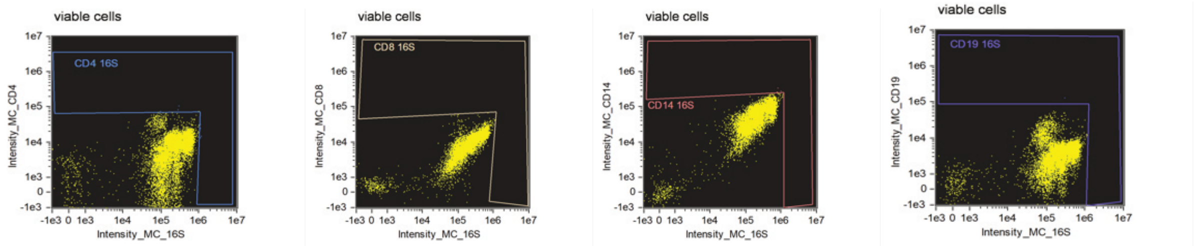


Intensity_MC_16S, Intensity_MC_CD4

| Population | Count | %Gated |
|---|---|---|
| viable cells & cells & F | 8827 | 100 |
| CD4 16S & viable cells & ce... | 7 | 0.08 |

Intensity_MC_16S, Intensity_MC_CD8

| Population | Count | %Gated |
|---|---|---|
| viable cells & cells & F | 8827 | 100 |
| CD8 16S & viable cells & ce... | 2 | 0.02 |

Intensity_MC_16S, Intensity_MC_CD14

| Population | Count | %Gated |
|---|---|---|
| viable cells & cells & F | 8827 | 100 |
| CD14 16S & viable cells & c... | 3 | 0.03 |

Intensity_MC_16S, Intensity_MC_CD19

| Population | Count | %Gated |
|---|---|---|
| viable cells & cells & F | 8827 | 100 |
| CD19 16S & viable cells & c... | 1 | 0.01 |

**B**

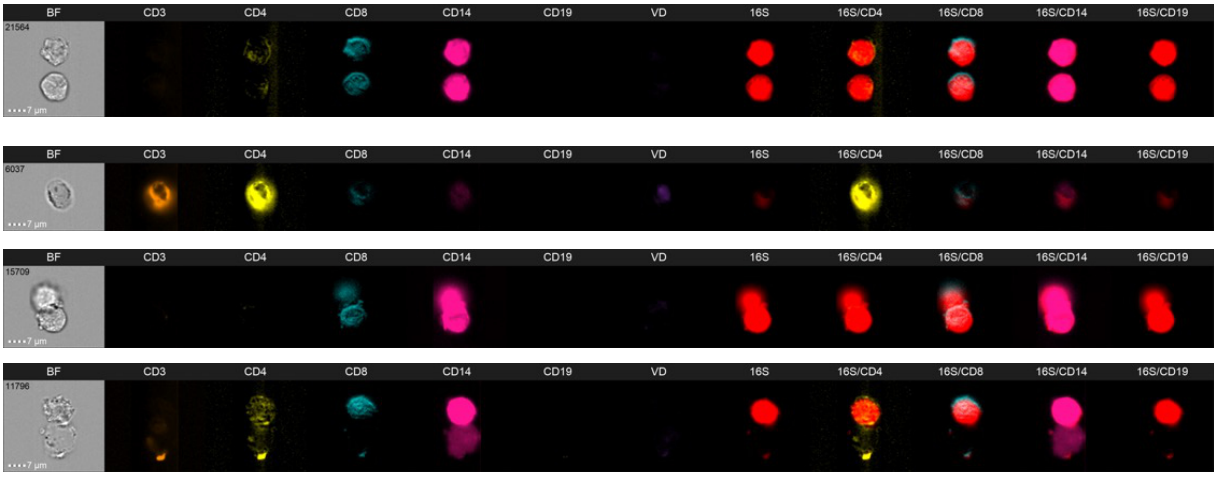

# S7

A

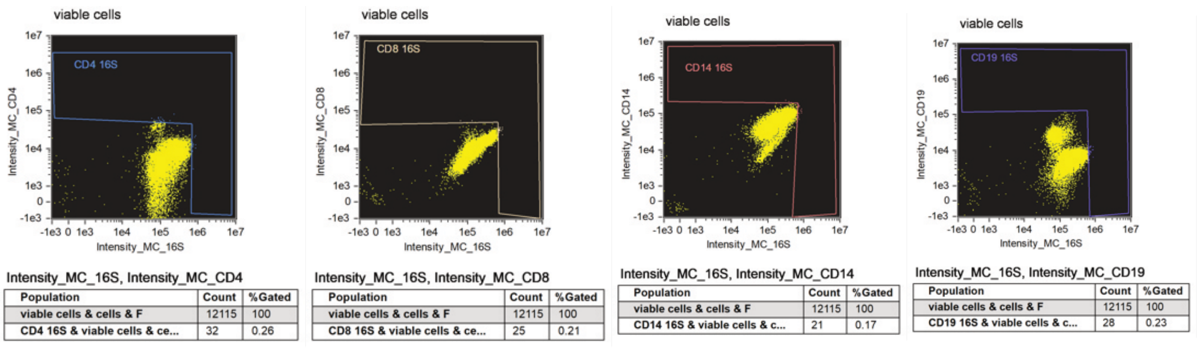
viable cells
CD4 16S
Intensity_MC_CD4
Intensity_MC_16S
Intensity_MC_16S, Intensity_MC_CD4
Population | Count | %Gated
viable cells & cells & F | 12115 | 100
CD4 16S & viable cells & ce... | 32 | 0.26
viable cells
CD8 16S
Intensity_MC_CD8
Intensity_MC_16S
Intensity_MC_16S, Intensity_MC_CD8
Population | Count | %Gated
viable cells & cells & F | 12115 | 100
CD8 16S & viable cells & ce... | 25 | 0.21
viable cells
CD14 16S
Intensity_MC_CD14
Intensity_MC_16S
Intensity_MC_16S, Intensity_MC_CD14
Population | Count | %Gated
viable cells & cells & F | 12115 | 100
CD14 16S & viable cells & c... | 21 | 0.17
viable cells
CD19 16S
Intensity_MC_CD19
Intensity_MC_16S
Intensity_MC_16S, Intensity_MC_CD19
Population | Count | %Gated
viable cells & cells & F | 12115 | 100
CD19 16S & viable cells & c... | 28 | 0.23

B

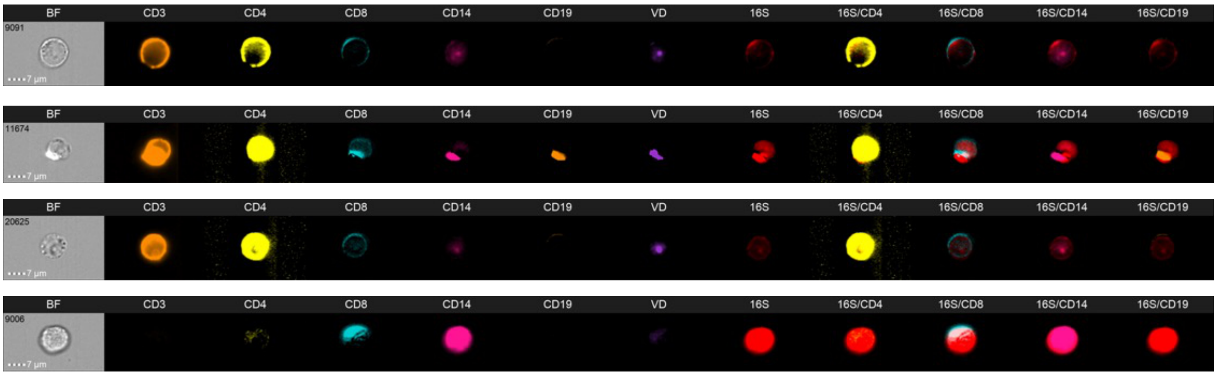
BF
CD3
CD4
CD8
CD14
CD19
VD
16S
16S/CD4
16S/CD8
16S/CD14
16S/CD19
9091
11674
20625
9006
7 µm

S8

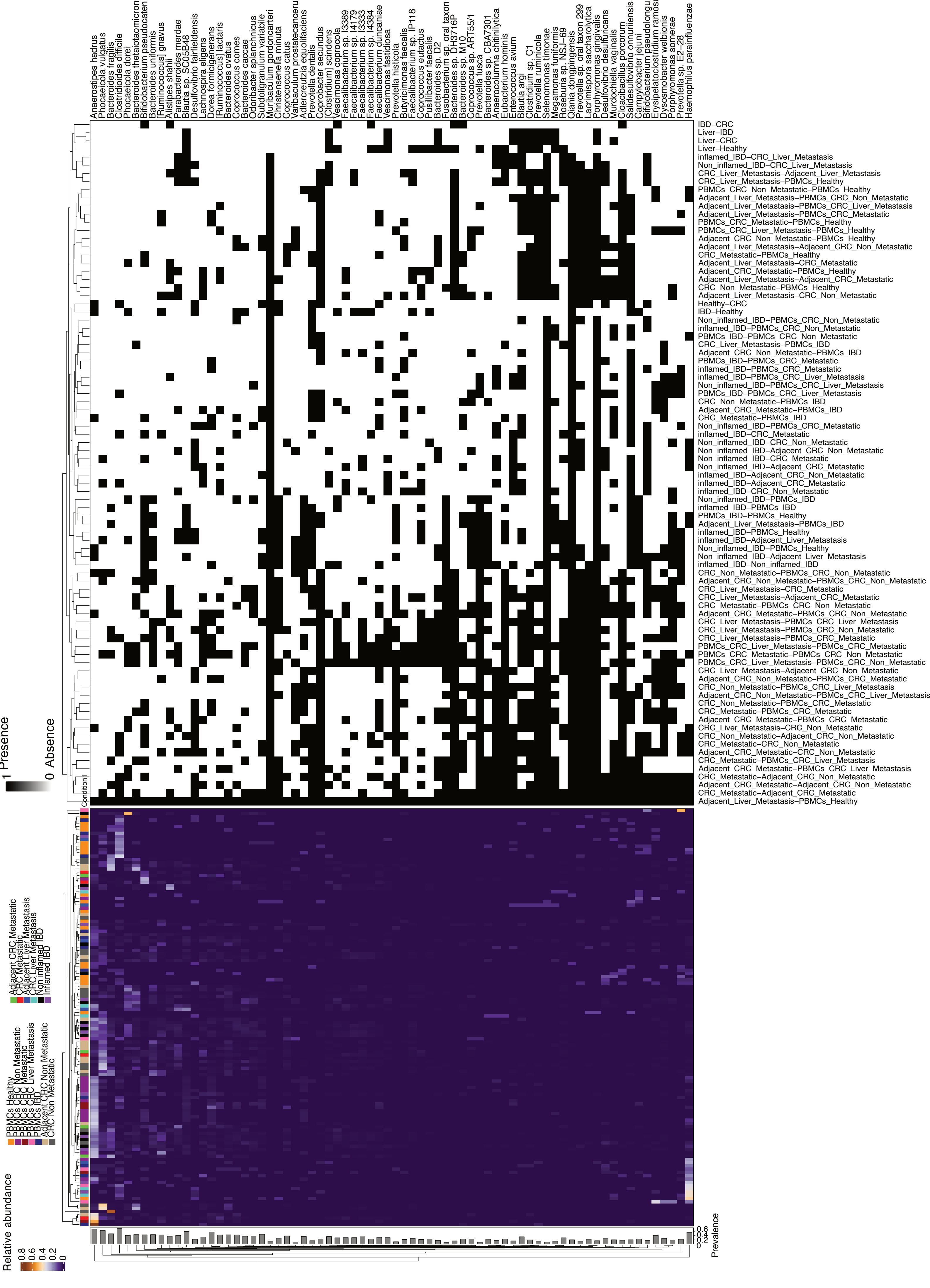
Anaerostipes hadrus
Phocaeicola vulgatus
Bacteroides fragilis
Clostridioides difficile
Phocaeicola dorei
Bacteroides thetaiotaomicron
Bifidobacterium pseudocatenulatum
Bacteroides uniformis
[Ruminococcus] gnavus
Alistipes shahii
Parabacteroides merdae
Blautia sp. SC05B48
Desulfovibrio fairfieldensis
Lachnospira eligens
Dorea formicigenerans
[Ruminococcus] lactaris
Bacteroides ovatus
Coprococcus comes
Bacteroides caccae
Odoribacter splanchnicus
Subdoligranulum variabile
Muribaculum gordoncarteri
Christensenella minuta
Coprococcus catus
Varibaculum prostatecancerukia
Adlercreutzia equolifaciens
Prevotella dentalis
Coprobacter secundus
[Clostridium] scindens
Vescimonas coprocola
Faecalibacterium sp. I3389
Faecalibacterium sp. I4179
Faecalibacterium sp. I3333
Faecalibacterium sp. I4384
Faecalibacterium duncaniae
Vescimonas fastidiosa
Prevotella histicola
Butyricimonas faecalis
Faecalibacterium sp. IP118
Coprococcus eutactus
Pusillibacter faecalis
Bacteroides sp. D2
Fusobacterium sp. oral taxon 203
Bacteroides sp. DH3716P
Bacteroides sp. M10
Coprococcus sp. ART55/1
Prevotella fusca
Bacteroides sp. CBA7301
Anaerocolumna chitinilytica
Eubacterium hominis
Enterococcus avium
Blautia argi
Clostridium sp. C1
Prevotella ruminicola
Selenomonas timonae
Megamonas funiformis
Roseburia sp. NSJ–69
Qiania dongpingensis
Prevotella sp. oral taxon 299
Lacrimispora saccharolytica
Porphyromonas gingivalis
Desulfovibrio desulfuricans
Murdochiella vaginalis
Cloacibacillus porcorum
Salidesulfovibrio onnuriiensis
Campylobacter jejuni
Bifidobacterium pseudolongum
Erysipelatoclostridium ramosum
Dysosmobacter welbionis
Porphyromonas somerae
Prevotella sp. E2–28
Haemophilus parainfluenzae
IBD–CRC
Liver–IBD
Liver–CRC
Liver–Healthy
inflamed_IBD–CRC_Liver_Metastasis
Non_inflamed_IBD–CRC_Liver_Metastasis
CRC_Liver_Metastasis–Adjacent_Liver_Metastasis
CRC_Liver_Metastasis–PBMCs_Healthy
PBMCs_CRC_Non_Metastatic–PBMCs_Healthy
Adjacent_Liver_Metastasis–PBMCs_CRC_Non_Metastatic
Adjacent_Liver_Metastasis–PBMCs_CRC_Liver_Metastasis
Adjacent_Liver_Metastasis–PBMCs_CRC_Metastatic
PBMCs_CRC_Metastatic–PBMCs_Healthy
PBMCs_CRC_Liver_Metastasis–PBMCs_Healthy
Adjacent_CRC_Non_Metastatic–PBMCs_Healthy
Adjacent_Liver_Metastasis–Adjacent_CRC_Non_Metastatic
CRC_Metastatic–PBMCs_Healthy
Adjacent_Liver_Metastasis–CRC_Metastatic
Adjacent_CRC_Metastatic–PBMCs_Healthy
Adjacent_Liver_Metastasis–Adjacent_CRC_Metastatic
CRC_Non_Metastatic–PBMCs_Healthy
Adjacent_Liver_Metastasis–CRC_Non_Metastatic
Healthy–CRC
IBD–Healthy
Non_inflamed_IBD–PBMCs_CRC_Non_Metastatic
inflamed_IBD–PBMCs_CRC_Non_Metastatic
PBMCs_IBD–PBMCs_CRC_Non_Metastatic
CRC_Liver_Metastasis–PBMCs_IBD
Adjacent_CRC_Non_Metastatic–PBMCs_IBD
PBMCs_IBD–PBMCs_CRC_Metastatic
inflamed_IBD–PBMCs_CRC_Metastatic
inflamed_IBD–PBMCs_CRC_Liver_Metastasis
Non_inflamed_IBD–PBMCs_CRC_Liver_Metastasis
PBMCs_IBD–PBMCs_CRC_Liver_Metastasis
CRC_Non_Metastatic–PBMCs_IBD
Adjacent_CRC_Metastatic–PBMCs_IBD
CRC_Metastatic–PBMCs_IBD
Non_inflamed_IBD–PBMCs_CRC_Metastatic
inflamed_IBD–CRC_Metastatic
Non_inflamed_IBD–CRC_Non_Metastatic
Non_inflamed_IBD–Adjacent_CRC_Non_Metastatic
Non_inflamed_IBD–CRC_Metastatic
Non_inflamed_IBD–Adjacent_CRC_Metastatic
inflamed_IBD–Adjacent_CRC_Non_Metastatic
inflamed_IBD–Adjacent_CRC_Metastatic
inflamed_IBD–CRC_Non_Metastatic
Non_inflamed_IBD–PBMCs_IBD
inflamed_IBD–PBMCs_IBD
PBMCs_IBD–PBMCs_Healthy
Adjacent_Liver_Metastasis–PBMCs_IBD
inflamed_IBD–PBMCs_Healthy
inflamed_IBD–Adjacent_Liver_Metastasis
Non_inflamed_IBD–PBMCs_Healthy
Non_inflamed_IBD–Adjacent_Liver_Metastasis
inflamed_IBD–Non_inflamed_IBD
CRC_Non_Metastatic–PBMCs_CRC_Non_Metastatic
Adjacent_CRC_Non_Metastatic–PBMCs_CRC_Non_Metastatic
CRC_Liver_Metastasis–CRC_Metastatic
CRC_Liver_Metastasis–Adjacent_CRC_Metastatic
CRC_Metastatic–PBMCs_CRC_Non_Metastatic
Adjacent_CRC_Metastatic–PBMCs_CRC_Non_Metastatic
CRC_Liver_Metastasis–PBMCs_CRC_Liver_Metastasis
CRC_Liver_Metastasis–PBMCs_CRC_Non_Metastatic
CRC_Liver_Metastasis–PBMCs_CRC_Metastatic
PBMCs_CRC_Liver_Metastasis–PBMCs_CRC_Metastatic
PBMCs_CRC_Metastatic–PBMCs_CRC_Non_Metastatic
PBMCs_CRC_Liver_Metastasis–PBMCs_CRC_Non_Metastatic
CRC_Liver_Metastasis–Adjacent_CRC_Non_Metastatic
Adjacent_CRC_Non_Metastatic–PBMCs_CRC_Metastatic
CRC_Non_Metastatic–PBMCs_CRC_Liver_Metastasis
Adjacent_CRC_Non_Metastatic–PBMCs_CRC_Liver_Metastasis
CRC_Non_Metastatic–PBMCs_CRC_Metastatic
CRC_Metastatic–PBMCs_CRC_Metastatic
Adjacent_CRC_Metastatic–PBMCs_CRC_Metastatic
CRC_Liver_Metastasis–CRC_Non_Metastatic
CRC_Non_Metastatic–Adjacent_CRC_Non_Metastatic
CRC_Metastatic–CRC_Non_Metastatic
Adjacent_CRC_Metastatic–CRC_Non_Metastatic
CRC_Metastatic–PBMCs_CRC_Liver_Metastasis
Adjacent_CRC_Metastatic–PBMCs_CRC_Liver_Metastasis
CRC_Metastatic–Adjacent_CRC_Non_Metastatic
Adjacent_CRC_Metastatic–Adjacent_CRC_Non_Metastatic
CRC_Metastatic–Adjacent_CRC_Metastatic
Adjacent_Liver_Metastasis–PBMCs_Healthy
1 Presence
0 Absence
Condition
Adjacent CRC Metastatic
CRC Metastatic
Adjacent Liver Metastasis
CRC Liver Metastasis
Non inflamed IBD
inflamed IBD
PBMCs Healthy
PBMCs CRC Non Metastatic
PBMCs CRC Metastatic
PBMCs CRC Liver Metastasis
PBMCs IBD
Adjacent CRC Non Metastatic
CRC Non Metastatic
Relative abundance
0.8
0.6
0.4
0.2
0
Prevalence
0.6
0.4
0.2
0

**S9**

**A** *Anaerostipes hadrus*

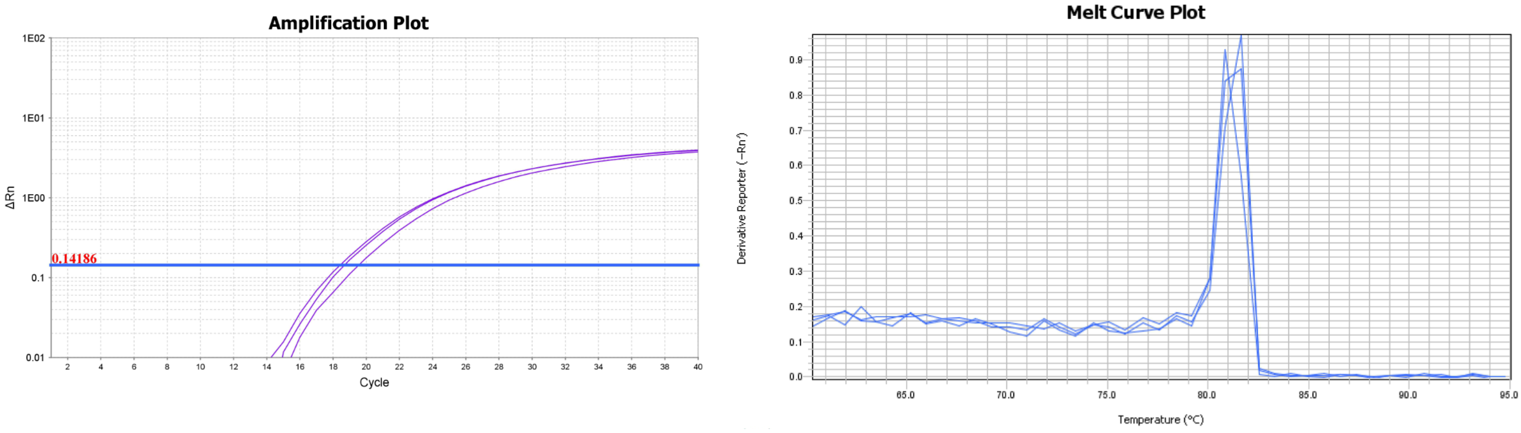


**B** *Bifidobacterium adolescentis*

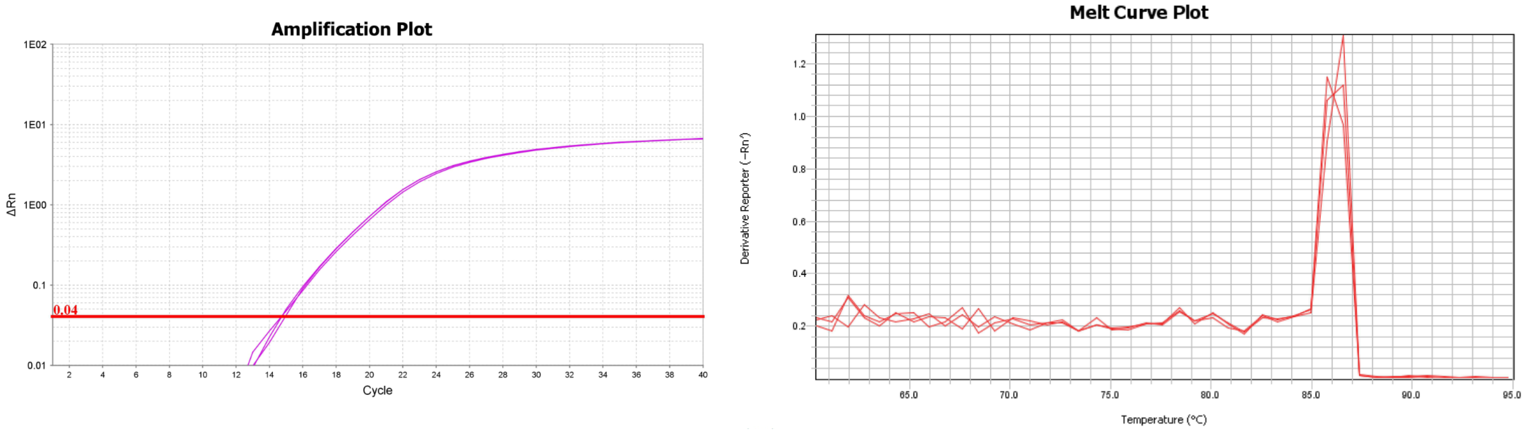


**C** *Collinsella aerofaciens*

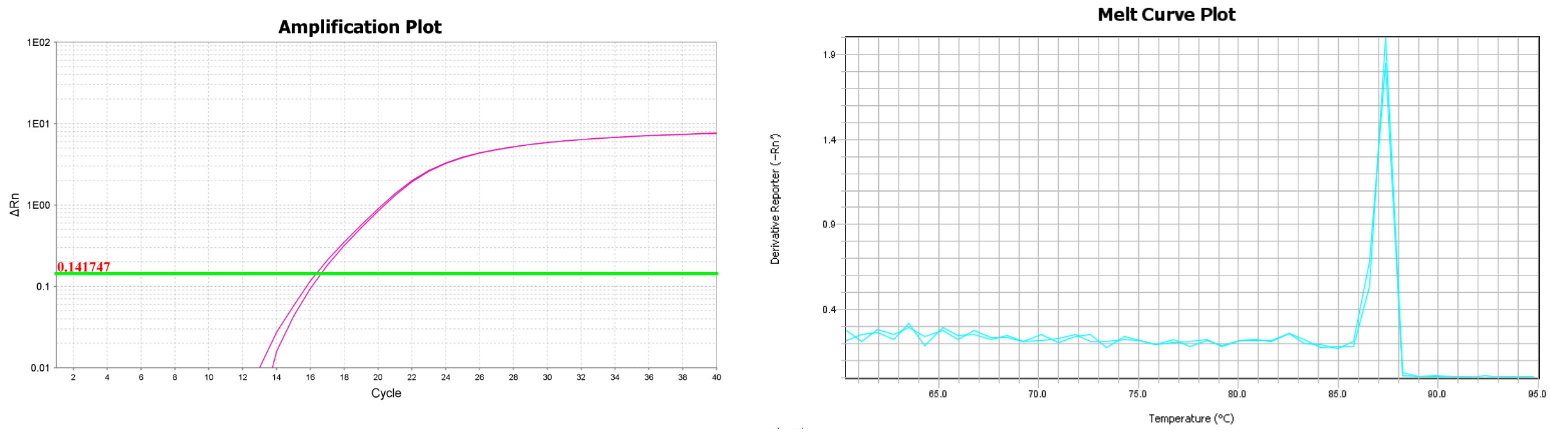


**D** *Faecalibacterium prausnitzii*

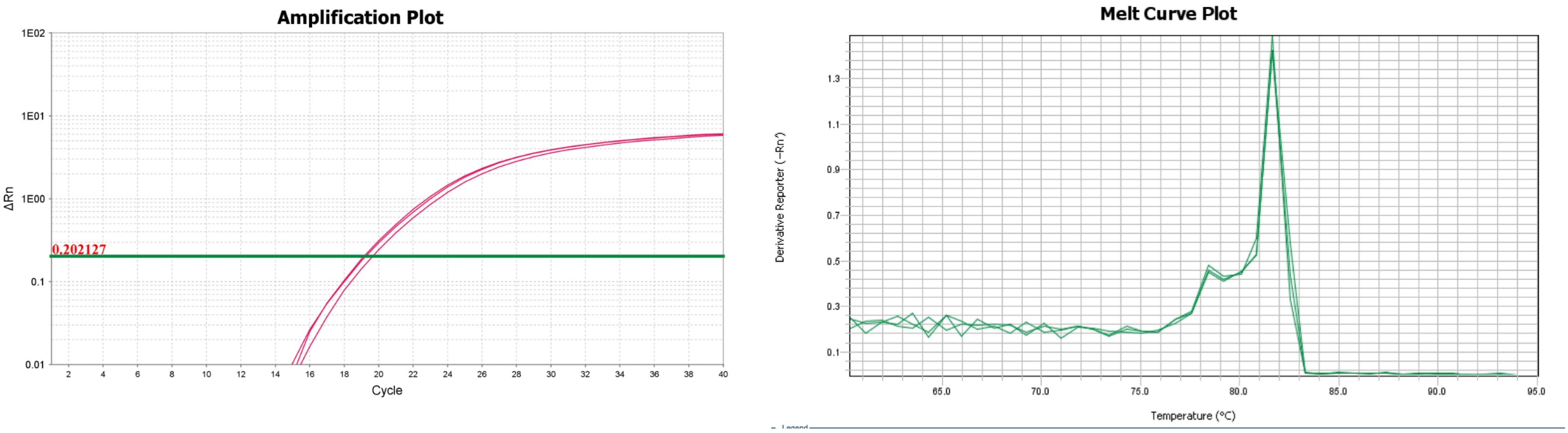

**S10**

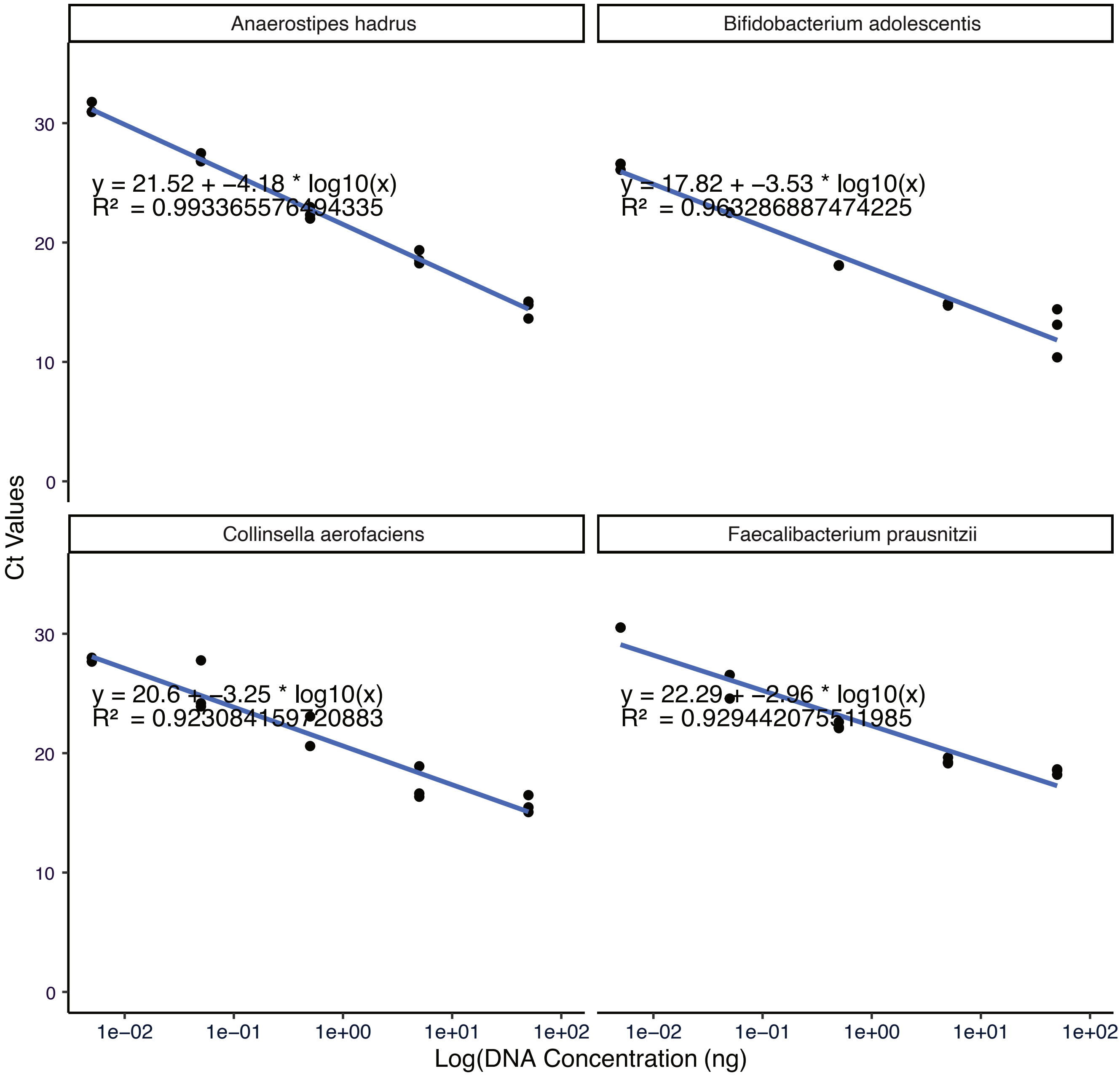

Anaerostipes hadrus
Bifidobacterium adolescentis
Collinsella aerofaciens
Faecalibacterium prausnitzii
y = 21.52 + −4.18 * log10(x)
R² = 0.993365576494335
y = 17.82 + −3.53 * log10(x)
R² = 0.963286887474225
y = 20.6 + −3.25 * log10(x)
R² = 0.923084159720883
y = 22.29 + −2.96 * log10(x)
R² = 0.929442075511985
Ct Values
0
10
20
30
1e−02
1e−01
1e+00
1e+01
1e+02
Log(DNA Concentration (ng)

**S11**

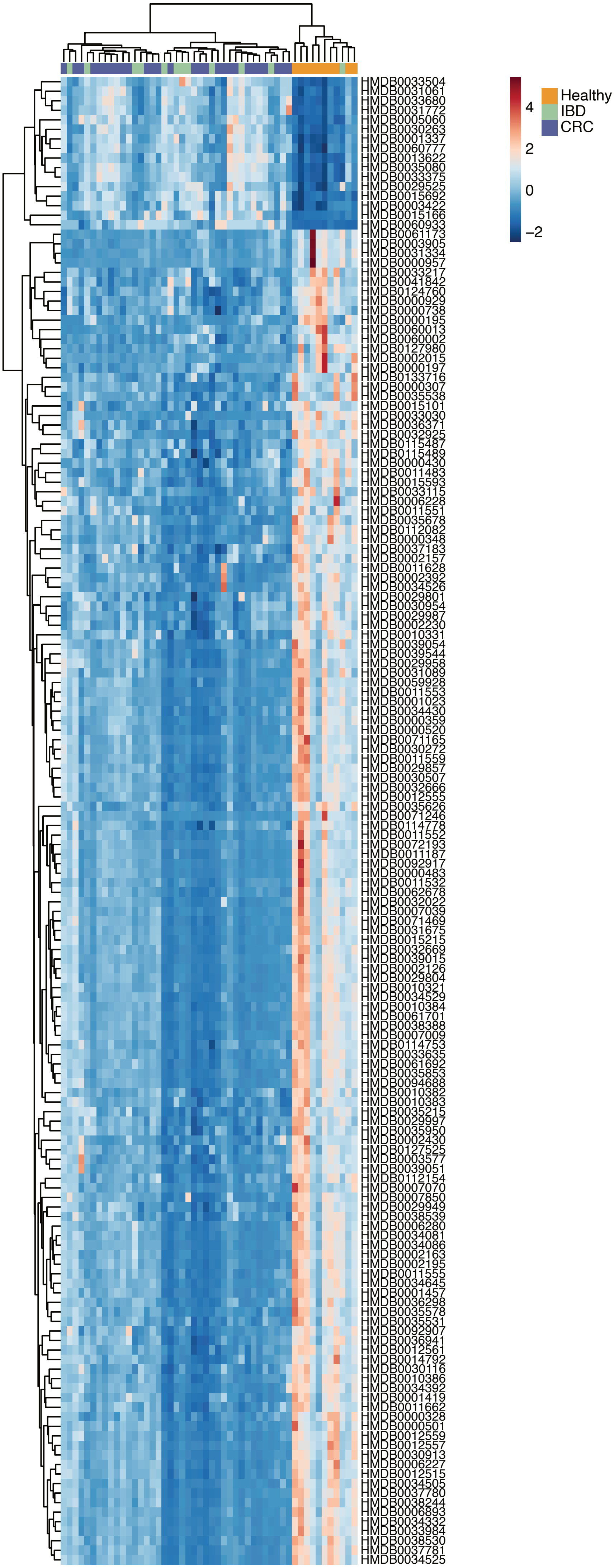
Healthy
IBD
CRC
4
2
0
−2
HMDB0033504
HMDB0031061
HMDB0033680
HMDB0031772
HMDB0005060
HMDB0030263
HMDB0001337
HMDB0060777
HMDB0013622
HMDB0035080
HMDB0033375
HMDB0029525
HMDB0015692
HMDB0003422
HMDB0015166
HMDB0060933
HMDB0061173
HMDB0003905
HMDB0031334
HMDB0000957
HMDB0033217
HMDB0041842
HMDB0124760
HMDB0000929
HMDB0000738
HMDB0000195
HMDB0060013
HMDB0060002
HMDB0127980
HMDB0002015
HMDB0000197
HMDB0133716
HMDB0000307
HMDB0035538
HMDB0015101
HMDB0033030
HMDB0036371
HMDB0032925
HMDB0115487
HMDB0115489
HMDB0000430
HMDB0011483
HMDB0015593
HMDB0033115
HMDB0006228
HMDB0011551
HMDB0035678
HMDB0112082
HMDB0000348
HMDB0037183
HMDB0002157
HMDB0011628
HMDB0002392
HMDB0034526
HMDB0029801
HMDB0030954
HMDB0029987
HMDB0002230
HMDB0010331
HMDB0039054
HMDB0039544
HMDB0029958
HMDB0031089
HMDB0059928
HMDB0011553
HMDB0001023
HMDB0034430
HMDB0000359
HMDB0000520
HMDB0071165
HMDB0030272
HMDB0011559
HMDB0029857
HMDB0030507
HMDB0032666
HMDB0012555
HMDB0035626
HMDB0071246
HMDB0114778
HMDB0011552
HMDB0072193
HMDB0011187
HMDB0092917
HMDB0000483
HMDB0011532
HMDB0062678
HMDB0032022
HMDB0007039
HMDB0071469
HMDB0031675
HMDB0015215
HMDB0032669
HMDB0039015
HMDB0002126
HMDB0029804
HMDB0010321
HMDB0034529
HMDB0010384
HMDB0061701
HMDB0038388
HMDB0007009
HMDB0114753
HMDB0033635
HMDB0061692
HMDB0035853
HMDB0094688
HMDB0010382
HMDB0010383
HMDB0035215
HMDB0029997
HMDB0035950
HMDB0002430
HMDB0127525
HMDB0003577
HMDB0039051
HMDB0112154
HMDB0007070
HMDB0007850
HMDB0029949
HMDB0038539
HMDB0006280
HMDB0034081
HMDB0034086
HMDB0002163
HMDB0002195
HMDB0011555
HMDB0034645
HMDB0001457
HMDB0036298
HMDB0035578
HMDB0035531
HMDB0092907
HMDB0036941
HMDB0012561
HMDB0014792
HMDB0030116
HMDB0010386
HMDB0034392
HMDB0001419
HMDB0011662
HMDB0000328
HMDB0000501
HMDB0012559
HMDB0012557
HMDB0030913
HMDB0006227
HMDB0012515
HMDB0034505
HMDB0037780
HMDB0038244
HMDB0006893
HMDB0034332
HMDB0033984
HMDB0038530
HMDB0037781
HMDB0034525

**S12**

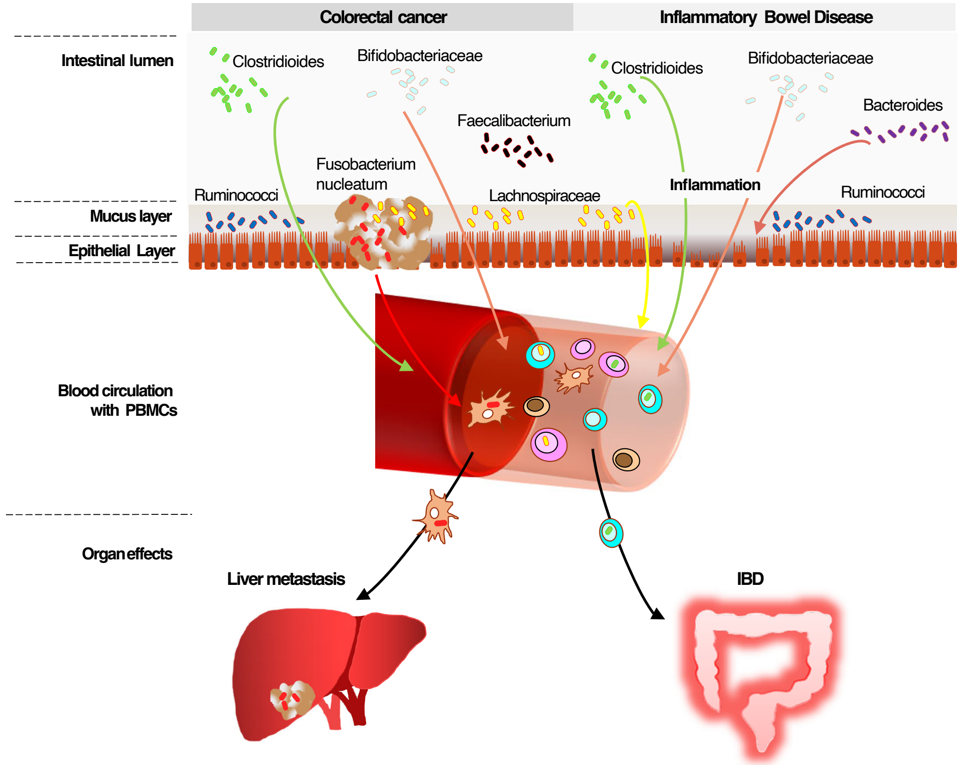
Colorectal cancer
Inflammatory Bowel Disease
Intestinal lumen
Clostridioides
Bifidobacteriaceae
Clostridioides
Bifidobacteriaceae
Faecalibacterium
Bacteroides
Fusobacterium nucleatum
Inflammation
Ruminococci
Lachnospiraceae
Ruminococci
Mucus layer
Epithelial Layer
Blood circulation with PBMCs
Organ effects
Liver metastasis
IBD
